# No evidence for a relationship between social closeness and similarity in resting-state functional brain connectivity in schoolchildren

**DOI:** 10.1101/788208

**Authors:** Carolyn Beth McNabb, Laura Grace Burgess, Amy Fancourt, Nancy Mulligan, Lily FitzGibbon, Patricia Riddell, Kou Murayama

**Affiliations:** School of Psychology and Clinical Language Sciences, University of Reading, Reading, United Kingdom RG6 7BE; BrainCanDo, Queen Anne’s School, Reading, United Kingdom RG4 6DX; Research Institute, Kochi University of Technology, Kami, Kochi, Japan 782-8502

**Keywords:** Adolescence, dyad, homophily, magnetic resonance imaging, social network

## Abstract

Previous research suggests that the proximity of individuals in a social network predicts how similarly their brains respond to naturalistic stimuli. However, the relationship between social connectedness and brain connectivity in the absence of external stimuli has not been examined. To investigate whether neural homophily between friends exists at rest we collected resting-state functional magnetic resonance imaging (fMRI) data from 68 school-aged girls, along with social network information from all pupils in their year groups (total 5,066 social dyads). Participants were asked to rate the amount of time they voluntarily spent with each person in their year group, and directed social network matrices and community structure were then determined from these data. No statistically significant relationships between social distance, community homogeneity and similarity of global-level resting-state connectivity were observed. Nor were we able to predict social distance using a machine learning technique (i.e. elastic net regression based on the local-level similarities in resting-state whole-brain connectivity between participants). Although neural homophily between friends exists when viewing naturalistic stimuli, this finding did not extend to functional connectivity at rest in our population. Instead, resting-state connectivity may be less susceptible to the influences of a person’s social environment.

---

Homophily is the tendency of individuals to attract and interact with those who share similar traits. Homophilic selection is observed for broad categorical traits such as gender, ethnicity and sexual orientation ^1-3^ but also for personal traits such as motivation ^4^, personality and cognitive ability ^5^, and academic achievement ^6^. High school and university students have been found to rearrange their local social networks to form ties and clusters with students who have similar performance levels ^6^ and this type of homophily has been observed even in polygenic scores for academic achievement ^7^.

Given the predominance of social network homophily for behavioural, personality and cognitive traits, we can reasonably expect that this extends to similarities in brain function. In fact, neural responses observed during unconstrained viewing of naturalistic stimuli (movie clips) were found to be significantly more similar among friends compared with those farther removed in a real-world social network ^8^. This effect persisted, even after controlling for inter-subject similarities in demographic variables, such as age, gender, nationality and ethnicity. Social closeness also provides opportunities for behavioural contagion - researchers have shown that social contagion modulates neural representations of risk in reward-related areas and that functional connectivity between the caudate and prefrontal cortex accounts for individual differences in susceptibility to risk-taking contagion ^9^.

Previously, neural similarity was assessed using intersubject correlation of blood oxygenation level-dependent (BOLD) timeseries across functionally derived regions of the brain. This method of inter-subject correlation evaluates the externally generated (extrinsic) stimulus-locked BOLD activation associated with the task but ignores the internally generated (intrinsic) component of BOLD activity, which is cancelled out when correlating across participants ^10^. Therefore, it remains unclear whether internally-generated brain activity similarly exhibits neural homophily between friends.

Patterns of brain connectivity elicited from internally generated resting-state BOLD activation are mirrored by activation networks found under explicit task-based activation ^11^. For example, resting-state sub-networks have been shown to correspond with externally generated activation from attention, speech, reasoning, emotion, memory and social cognition tasks ^11,12^. Resting-state connectivity is also associated with non-cognitive measures of motivation. Grit and growth mind-set were found to be associated with functional connectivity between ventral striatal and bilateral prefrontal networks important for cognitive-behavioural control ^13^. Connectivity at rest also predicts personality. Connectome-based predictive modelling has been used to successfully predict trait-level measures of personality, including openness to experience ^14^, neuroticism and extraversion ^15^. Others have found that global connectivity of the left prefrontal cortex predicts individual differences in fluid intelligence and cognitive control ^16^ and a clinical measure of attention can be predicted from resting-state connectivity in a network associated with sustained attention ^17^. These findings highlight the utility of resting-state connectivity for identifying individual differences in cognition, behaviour and personality, all of which have exhibited homophily within social networks.

Researchers have also linked internally generated brain connectivity with a number of social behaviours. For example, resting-state sub-networks for motor, visual, speech and other language functions have been associated with the quality and quantity of social networks in older adults ^18^. Others have demonstrated positive associations between functional connectivity and social network size and embeddedness ^19^. There is also evidence for stronger amygdalar connectivity with brain networks subserving perceptual and affiliative behaviours in healthy adults who foster and maintain larger and more complex social networks ^20^. Social network size may also dictate the degree of connectivity within the default mode network (DMN). The DMN overlaps considerably with regions important for theory of mind and social cognition ^12,21^ and has been found to exhibit greater coupling with anterior cingulate and dorsolateral prefrontal cortex in those with larger social network size ^22^. This striking overlap between the DMN and regions involved in social cognition infers a tendency for entertaining thoughts about oneself and others during rest ^23^.

However, despite investigations into the relationship between internally generated connectivity patterns and social behaviour, no study has investigated whether close social relationships are associated with similarities in resting-state connectivity. If behaviours and personality traits exhibit homophily and these traits have connectivity signatures in the resting brain, it may be possible for resting-state brain connectivity to also exhibit homophily. Therefore, in the current study, we set out to investigate whether voluntarily spending large amounts of time with a peer is correlated with a higher degree of resting-state similarity compared with peers who voluntarily spend less or no time with one another. Secondary school offers an excellent environment for investigating such associations; young adolescents attending secondary school frequently spend large quantities of time with one another while forming new and long-lasting relationships with their peers. Therefore, opportunities for behavioural (and potentially neural) contagion and selection at this age are high.

In this study, we collected resting-state fMRI data of pupils attending a single high school, along with social network (friendship) information between them. These unique data allowed us to test the hypothesis that friends exhibit greater similarity in internally generated functional brain connectivity compared with those farther removed in a school-based social network.

As homophily has been observed for traits such as motivation ^4^, personality and cognitive ability ^5^, we hypothesised that social closeness would result in higher levels of between-subject correlations in resting-state networks related to cognitive performance, social interaction and motivational processing. Specifically, we chose to focus on similarities within the DMN ^24^, salience network ^25^, and left and right frontoparietal networks (lFPN and rFPN, respectively) ^26^. To account for potential similarities outside of these functional networks, we also sought to investigate similarities in whole-brain connectivity between friends, hypothesising that friends may select one another based on a range of neural attributes not constrained to a single cognitive network.

In addition to testing our main hypotheses, we conducted exploratory analyses to investigate whether friends exhibit similarities in brain network organisation and communication using graph-theoretical analysis. Measures of interest included modularity - how easily the brain can be divided into distinct functional networks, regional (nodal) strength - the importance of a region within its network, and nodal diversity - a measure of regional integration. Finally, these analyses were complimented by a machine learning approach to social distance prediction based on neural similarity.

## Results

### Participant and network characteristics

This study included a total of 175 schoolchildren (5,066 dyads) from a local day and boarding school, of whom 68 (767 dyads) participated in the fMRI study. Participants spanned across two academic years, including two year 8 cohorts (12-13 years of age) and one year 9 cohort (13-14 years of age). This resulted in 5,066 dyads in the social network study and 767 dyads in the fMRI study. Demographic data for each cohort are presented in Table 1. FMRI cohorts were relatively well matched to full year group cohorts for most demographic characteristics. The fMRI sample size was sufficient to detect a small effect with high power (i.e. 80% or 95%) based on an in-house simulation (see Supplementary Materials: Study Power).

**Table 1.**
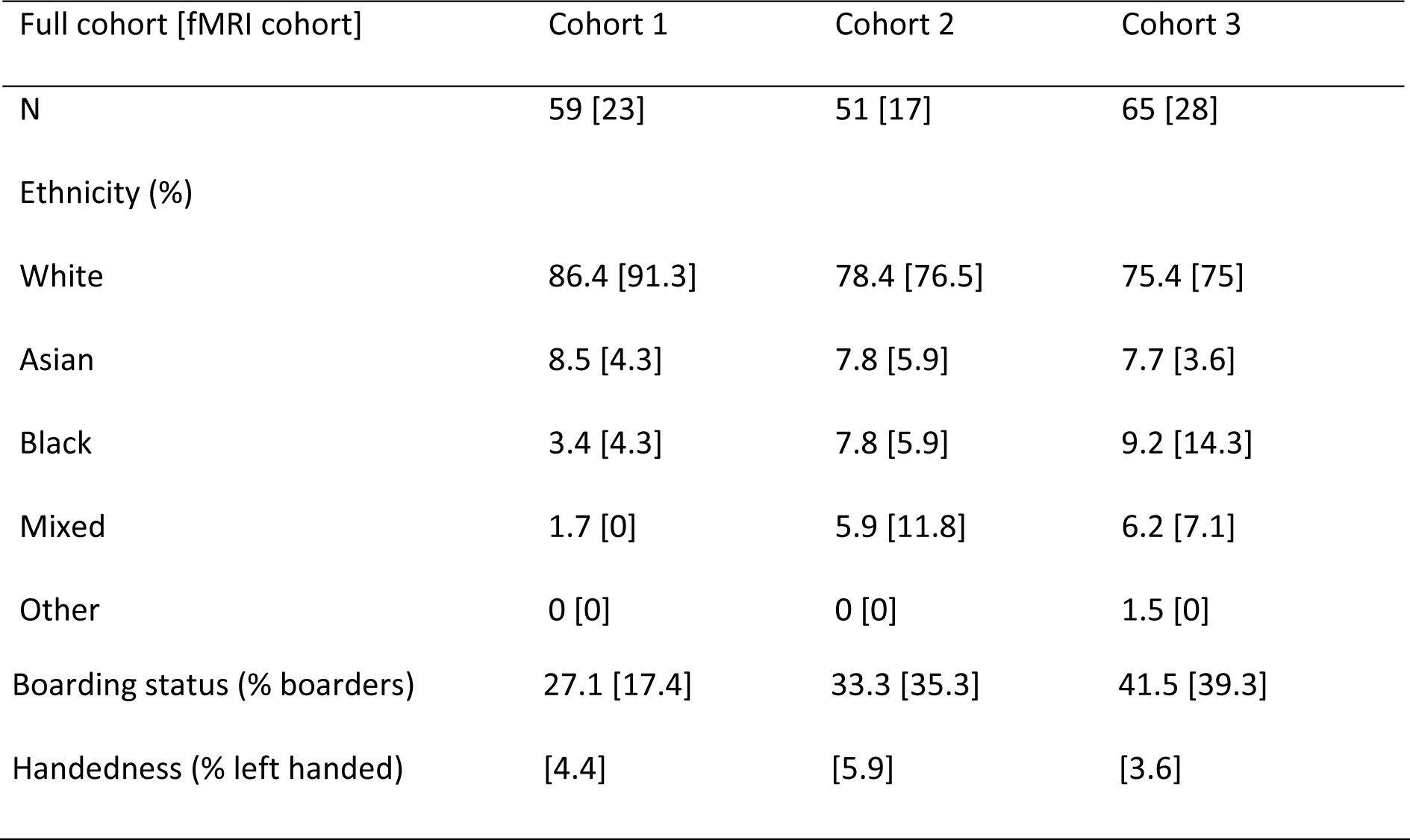
Demographic data for full and fMRI cohorts.

Participants from each cohort provided social network information (i.e. how much time they voluntarily spent with each person in their year group) for every other member of their cohort, providing almost complete networks of social connections for the three distinct cohorts of students. A roster-and-rating method was used. Specifically, participants were provided with a list of all students in their year group and asked to consider the question: “How much time do you spend interacting with this student?”. Students answered on a five-point Likert scale, which included options: “None”, “A rare amount”, “Some”, “More than some” and “Most”. Participants were told to consider time spent voluntarily interacting with other students but not time spent in planned seating situations, allocated group work or in classes without opportunities to talk amongst themselves.

In addition, participants were asked to nominate up to five individuals from their cohort with whom they considered themselves to be “close with” ^27^. Peer nominations and roster-and-rating methods measure different aspects of peer interactions. Whereas rating assesses general acceptance of peers, nomination is thought to encourage the naming of “best friends” ^28^. In the main text, we report results from roster-and-rating data only. However, nomination data provided the same conclusion as roster-and-rating data (see Supplementary Fig. S1 and S2).

Social networks were represented by unweighted, undirected, graphs. Social ties were only considered successful if participants rated spending “more than some” or “most” of their time with another student. Adjustment to include only ties where students spent “most” of their time together did not change the outcome of the results (see Supplementary Fig. S3 and S4). Mutually-reported (reciprocal) social ties are deemed to be more robust indicators of friendship than unreciprocated ties and were used by Parkinson et al. in their investigation of neural similarity during naturalistic viewing ^8^. For consistency with previous research, only reciprocal ties were included in the social network graphs, although use of non-reciprocated (directed) social ties did not change the overall results of the analysis (see Supplementary Fig. S5 and S6).

Each cohort was described in terms of its network characteristics, in particular, its network diameter, modularity, mean path length, reciprocity and density (see Supplementary Materials: Social Network Metrics for definitions of these measures). Network characteristics for each cohort are presented in Table 2. Relatively higher modularity and mean path length in cohort 3 suggest that this network was more segregated and less well integrated compared with the younger two cohorts (cohorts 1 and 2). The social networks for each cohort are depicted in Figure 1; fMRI cohorts were well distributed within whole year group samples and include both highly influential and less influential students (determined by Eigenvector centrality).

**Table 2.** Network characteristics of cohorts 1, 2 and 3, using threshold 4 social network data. Network diameter is the length of the longest geodesic distance between two nodes in the network. Modularity is a measure of how easily a network segregates into smaller subnetworks; large values represent networks that segregate easily into smaller communities. Mean path length is the mean geodesic distance between any two nodes in the network; smaller values are representative of more “tight-knit” networks. Reciprocity defines the proportion of connections in a directed graph that are mutual connections. Graph density gives the ratio of the number of connections (edges) and the number of possible connections in the network; higher values indicate that a larger number of possible connections have been made.

| Network characteristics | Cohort 1 | Cohort 2 | Cohort 3 |
| --- | --- | --- | --- |
| Network diameter | 3 | 3 | 4 |
| Modularity | 0.304 | 0.286 | 0.426 |
| Mean path length | 1.769 | 1.750 | 2.033 |
| Reciprocity | 0.618 | 0.630 | 0.668 |
| Graph density | 0.285 | 0.320 | 0.220 |

**Figure 1.**
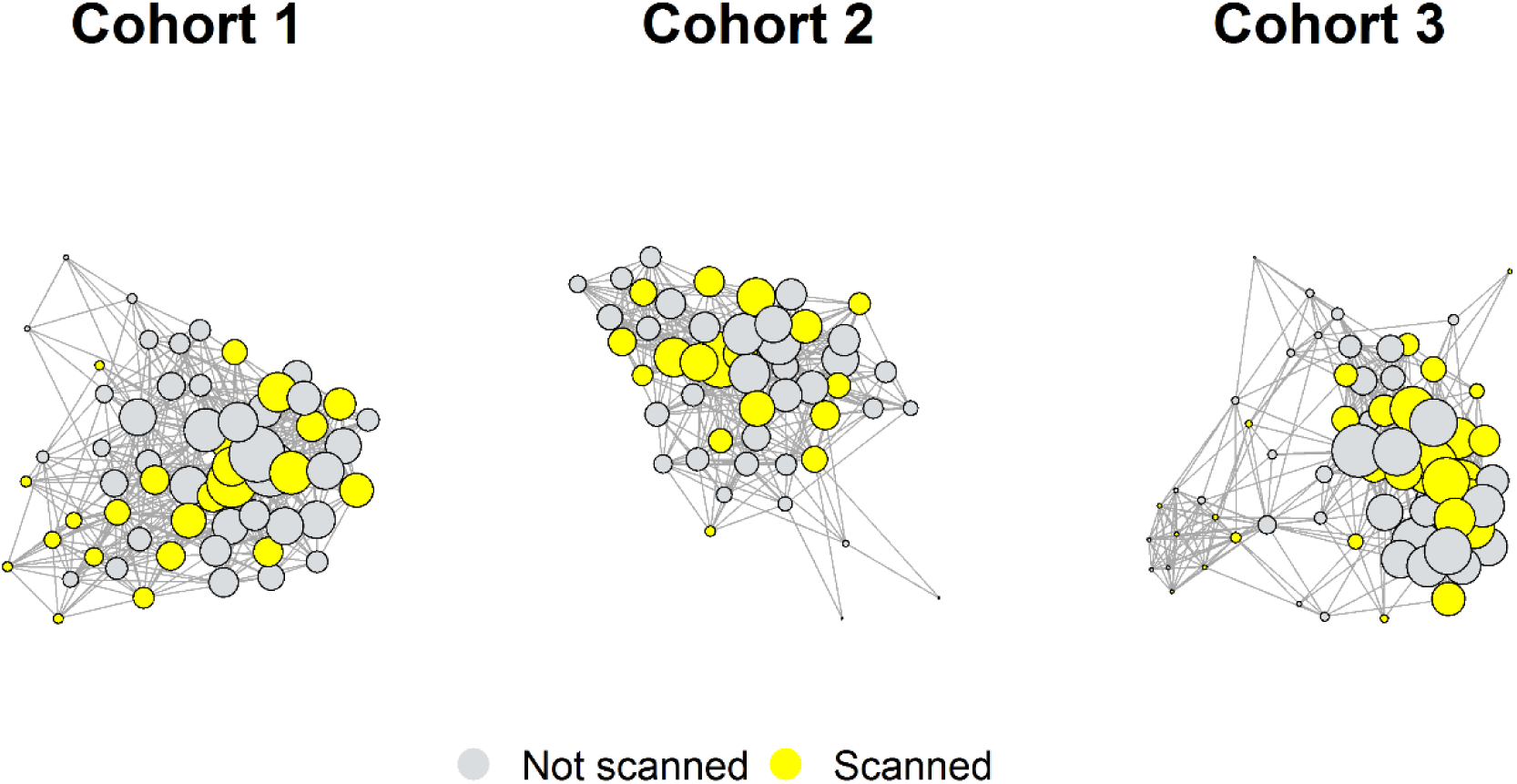
Social networks of cohorts 1, 2 and 3, depicting reciprocal friendships. Nodes represent students; lines (edges) represent mutually reported social ties where students rated the amount of time they spent with each other as “more than some” or “most” of the time. FMRI cohorts are depicted in green; students who provided information about their social interactions but were not included in the fMRI cohort are shown in grey. The size of each node depicts the Eigenvector centrality of that student. Eigenvector centrality is a measure of the relative importance/influence of a node in the network. Nodes with high importance (those who are themselves well-connected and are connected to others who are well-connected) have higher centrality (these are the largest nodes in the network), those with low importance have low centrality (these are the smallest nodes in the network).

The distance between two individuals in a social network is an important predictor of behavioural tendencies ^29^ and may relate to shared patterns in brain function ^8^. Social distances in the current study were relatively short compared with previous literature (e.g. Parkinson et al. (2018) reported a network diameter of 6 using only mutually reported social ties compared to a maximum diameter of 4 in the current study - see Table 2). The highly interconnected nature of our networks may affect how social distance and brain function similarities are associated with one another. Therefore, we also evaluated dyadic similarities as a function of community affiliation. By splitting social networks into smaller friendship communities and evaluating the differences in brain similarity between those inside and outside each community, we measured relationships between neural homophily and social behaviour at a binary friendship group level.

Measures of social proximity (social distance and community affiliation) were determined separately for each cohort. Social distance was calculated as the shortest path length between each pair of nodes (students) in the network. A dyad (student pair) with a social distance of 1 represented a relationship in which both students had said they spent “more than some” or “most” of their time with the other student (i.e. they were friends). Social distances of 2 and 3 represented dyad pairs in which students did not possess a reciprocal friendship (i.e. did not have a mutual rating of “more than some” or “most” of the time) but reported a mutual friend or friend of a friend, respectively.

Community structure was ascertained using the Louvain method ^30^. This method implements multi-level modularity optimisation to subdivide the network into non-overlapping groups of nodes that maximise the number of within-group (within-module) friendships (edges) and minimised the number of between-group friendships for each cohort. Community detection analysis identified four communities (modules) in cohorts 1 and 3 and three communities in cohort 2 (Figure 2). These communities were used in later analyses to classify student dyads in a binary manner, as either sharing a community or coming from different communities.

**Figure 2.**
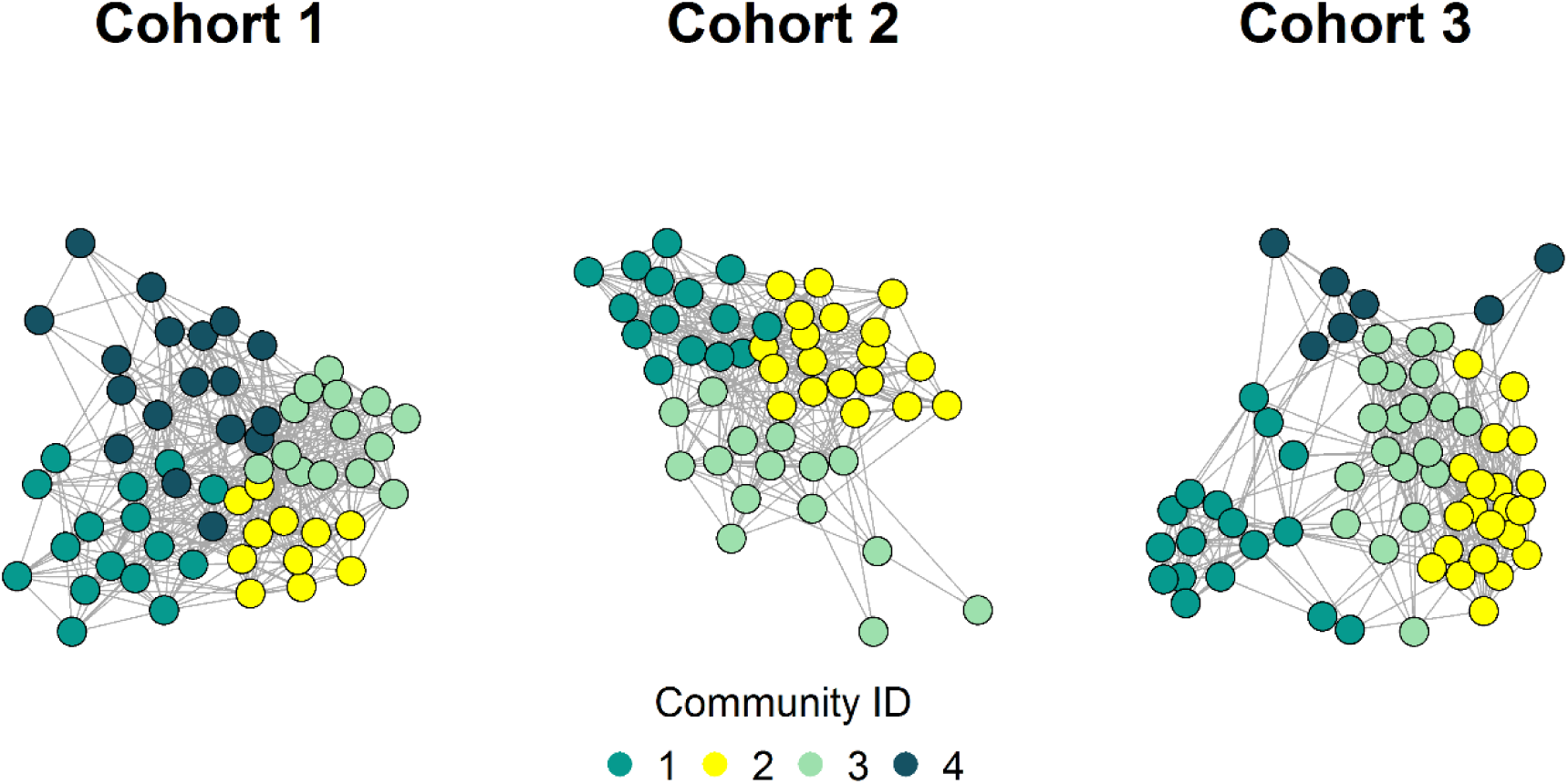
Community structure of each cohort, determined using the Louvain method ^30^. Community detection was used to estimate “friendship” groups within each cohort. The fMRI study included students from all communities.

### Main analyses: Dyadic similarities in functional connectivity as a function of social distance and community affiliation

A subset of participants (n=68, see Table 1 for details) participated in the fMRI component of the study. Functional echo-planar images were acquired over 10 minutes while participants rested with eyes open in the scanner (see Supplementary Materials for details of MRI acquisition). Mean blood oxygen level dependent (BOLD) time series were extracted from 272 anatomical regions of the brain for every participant in the fMRI cohort. Next, the Pearson’s correlation coefficient (connection strength) between BOLD time series was determined for each pair of regions (nodes) in MATLAB 2016a (MathWorks, USA). This produced a weighted, undirected whole-brain matrix of functional connectivity for each participant. *Z* transformations of weighted, undirected whole-brain matrices were then used for inter-subject correlations (Figure 3). Resting-state networks (DMN, salience, lFPN and rFPN) were defined based on previously published data (see Supplementary Materials: Resting-State Network Analysis). Nodes from the whole-brain parcellation that overlapped with at least 50 voxels from the resting-state networks were used to partition the whole-brain node-to-node matrix (see Supplementary Materials: **Error! Reference source not found.**). Data from each participant were compared with every other participant from the fMRI cohort using a Pearson’s correlation.

**Figure 3.**
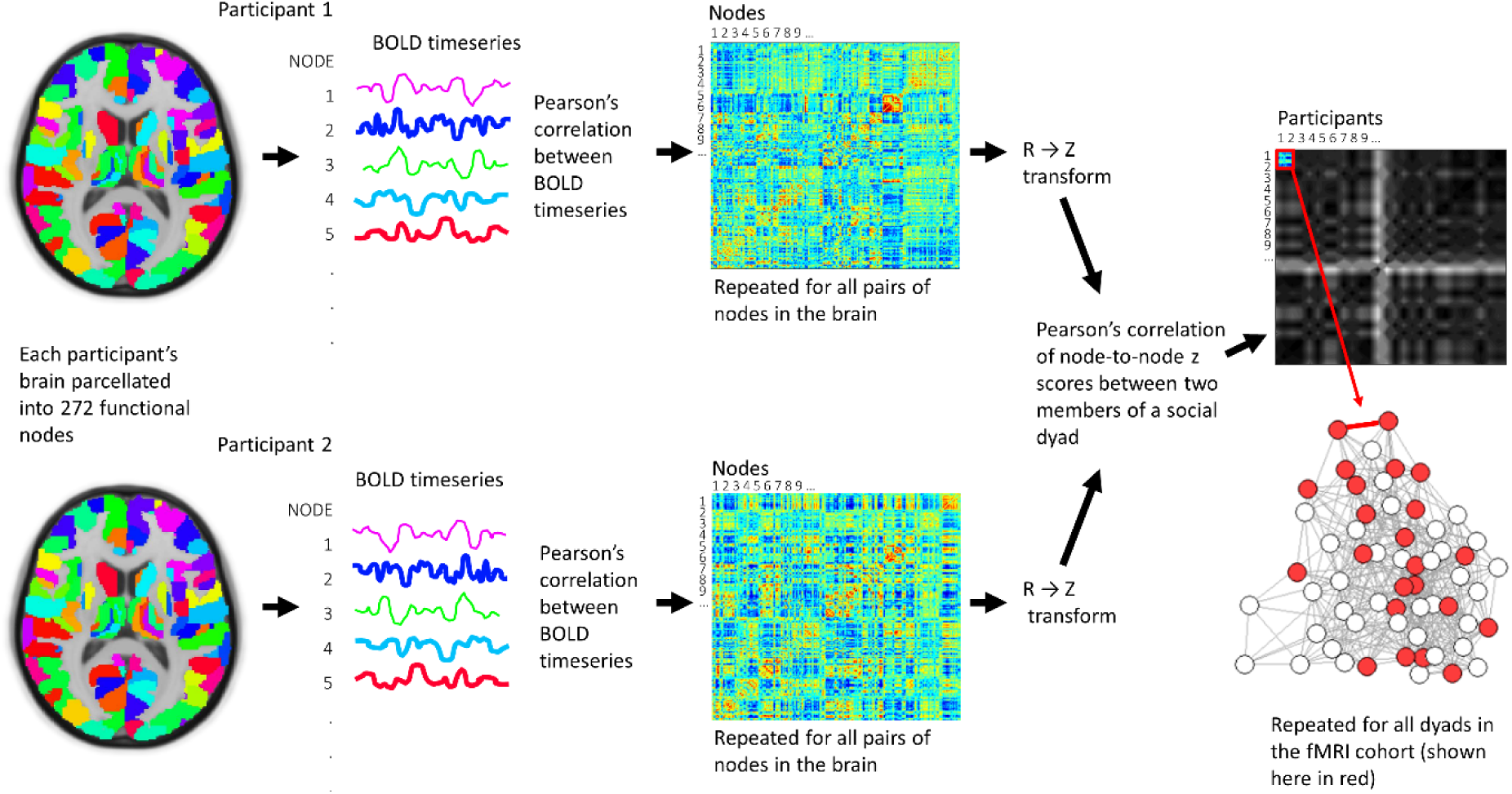
Processing pipeline for determining similarity of resting-state functional connectivity between participants. Mean blood oxygen level-dependent (BOLD) time series data for each brain parcel were correlated within participants and then each participant’s and then whole-brain correlation matrices were compared between participants.

Similarities in functional connectivity between participants were assessed using linear mixed effects modelling (LMEs) on a whole-brain network level as well as within individual resting-state networks for each fMRI cohort. Two LMEs (one for social distance and one for community affiliation) were applied to the data at each brain network level. Each student dyad (pair of students) in the fMRI cohort was described by one of two independent variables: social distance (i.e. shortest path length between the pair within the full year group cohort) or community affiliation (i.e. whether or not two students belonged to the same community, determined using the Louvain community detection method). The dependent variable for both models was similarity (measured as the correlation strength between two students) in resting-state connectivity, determined by the correlation strength of time series from each regional pair in the whole-brain or resting-state network (see Figure 3). Each student (*i* and *j*) in the dyad was modelled as a random effect to account for dependency introduced by the dyadic nature of the social network data.

No statistically significant relationship between social distance and similarity in functional brain connectivity was observed for any fMRI cohort at any resting-state network level (see Figure 4 for example plot). Beta weights (slopes) and 95% confidence intervals for LME models are presented in Figure 5a. Corresponding *t* and *p* values for individual tests are provided in the Supplementary Materials (Table S2). Cohorts were initially analysed separately to preserve the independence of the three social networks. We then integrated the regression coefficients of the LME models from three cohorts using a random-effect meta-analysis. Meta-analyses of LME models did not reveal any significant effects of social distance on the degree of functional brain similarity between students. These results indicate that the minimum path length between two individuals in a social network is not associated with similarities in brain function at rest, either at a whole-brain network or resting-state network level.

**Figure 4.**
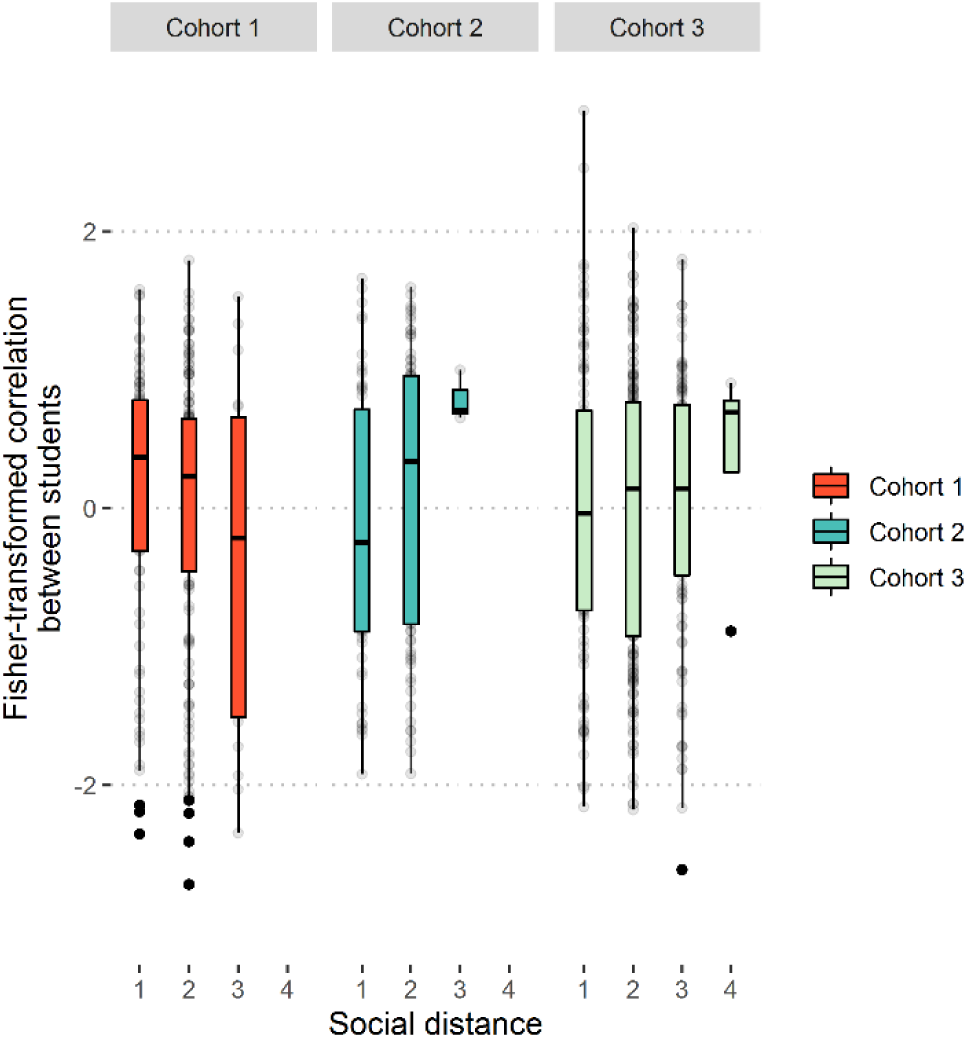
Example plot of standardised correlation strength of whole-brain connectivity between dyad members (y axis), as a function of social distance (x axis). Data here are for whole-brain connectivity similarities for all three cohorts; threshold was set at a distance of 4 (I spend “more than some” of my time with this person).

**Figure 5.**
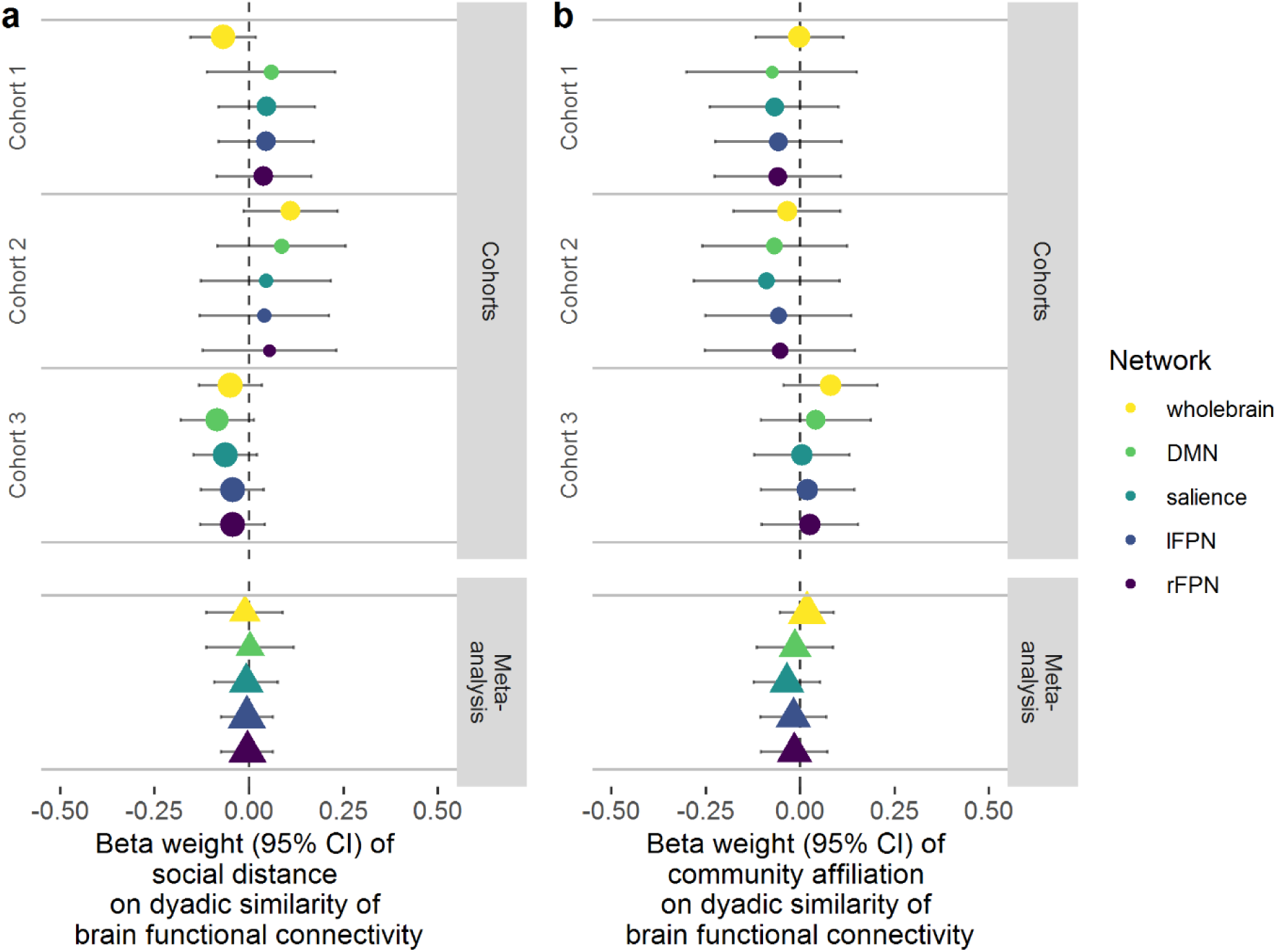
a) LME model outcomes and meta-analyses of functional connectivity similarity as a function of social distance. b) LME model outcomes and meta-analyses of functional connectivity similarity as a function of community affiliation. Whole-brain and resting state sub-network data are shown for all three cohorts. Circles and bars represent beta weights (slopes) and 95% confidence intervals (95% CI), respectively, for individual cohorts. Triangles represent beta weights for meta-analyses. Relative confidence in the effect is represented by the size of the circle/triangle; colours represent brain networks.

As our social networks were very densely connected, we opted to also represent dyadic relationships according to whether students belonged to the same “community”, determined by the Louvain community detection algorithm. Students’ relationships were defined by their affiliation to different communities (friendship groups), whereby dyads including students from the same community were scored 1 and dyads including students from different communities were scored 0. LME models predicting brain similarity as a function of community affiliation did not support an effect of friendship grouping as a predictor of connectivity similarity (i.e. the Pearson’s correlation strength between two individuals). This was true for whole-brain and resting-state network connectivity (Figure 5b). These results are consistent with the social distance analysis. They indicate that the similarity in resting brain function is no greater between individuals in the same social community (which we used here to estimate friendship groups) than those spanning different communities. Inclusion of demographic variables (i.e. similarities in ethnicity and boarding status) in LME models did not affect the statistical significance of the overall findings (see Table S3 for *t* and *p* values of individual tests).

### Exploratory analysis: graph theory measures of connectivity

Graph metrics of functional connectivity were derived from whole-brain weighted, undirected matrices using the Brain Connectivity Toolbox ^31^. Analysis steps are provided in Supplementary Materials: **Error! Reference source not found.**. Modularity and community structure were calculated using the Louvain method ^30^, assigning higher values to positively, compared with negatively weighted connections.

As for the main analyses, there were no statistically significant relationships between the similarity in strength, diversity or modularity of students’ brains and their distance from one another in the social network (Figure 6a). Nor were there any associations between these measures and the similarity in community affiliation (Figure 6b). These results add weight in support of the null hypothesis, that there is no relationship between social distance or community affiliation and similarity in resting-state brain connectivity.

**Figure 6.**
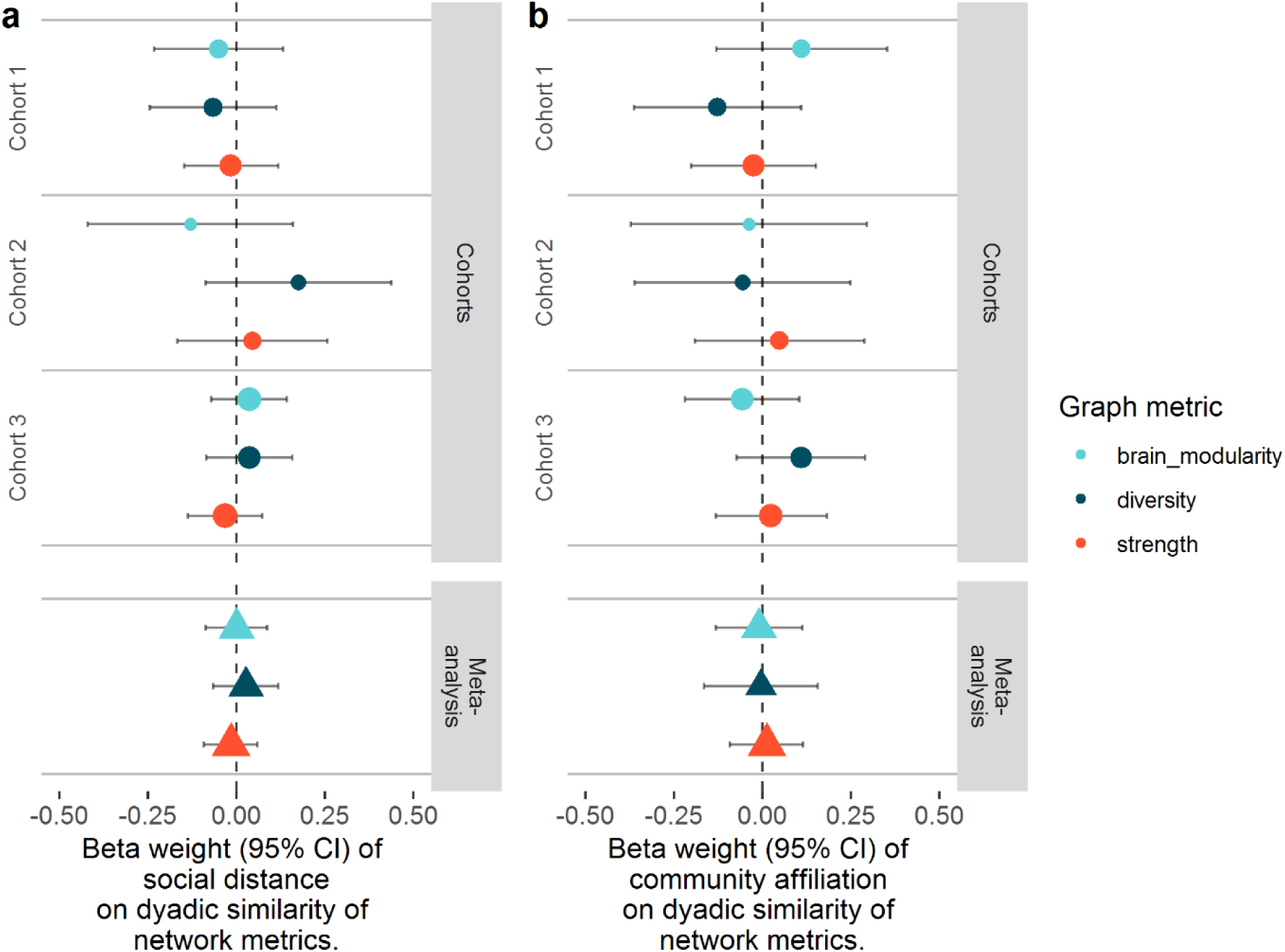
a) LME model outcomes and meta-analyses of similarities in graph theory measures of nodal strength, nodal diversity and brain modularity as a function of social distance. b) LME model outcomes and meta-analyses of similarities in graph theory measures of nodal strength, nodal diversity and brain modularity as a function of community affiliation. Data are shown for all three cohorts. Circles and bars represent beta weights (slopes) and 95% CI, respectively, for individual cohorts. Triangles represent beta weights for meta-analyses. Relative confidence in the effect is represented by the size of the circle/triangle; colours represent graph metrics.

### Exploratory analyses: Data-driven predictive model of social proximity from neural similarity

In the previous analyses, we assessed the overall similarity of the whole brain network or resting-state networks between pairs of students and examined whether social distance is related to the overall similarity. Although this analysis gives us the most straightforward test of our hypothesis, the analysis does not address the possibility that social distance is represented by the collection of local-level similarities (i.e. similarity between a specific pair of nodes). To examine whether any local-level similarity in the brain functional connectivity encodes social distance, we employed regularised (elastic net) regression techniques to predict social distance between two students based on similarities in their functional brain connectivity of all pairs of nodes. This regression technique has been used to successfully predict a variety of outcomes from MRI data, including openness to experience (a Big Five personality trait) ^14^, psychosis ^32^, and progression to Alzheimer’s disease in people with mild cognitive impairment ^33^, demonstrating its ability to cope with the high dimensionality of MRI data. In summary, predictive models performed poorly and failed to predict social distance with sufficient accuracy in any cohort. Further details of analysis and detailed results can be found in the Supplementary Materials. These data further support our main findings that resting state connectivity similarity between peers is not associated with social distance.

## Discussion

Our results provide little evidence for homophily of internally generated (resting-state) functional brain connectivity in school-based social networks. Neither whole-brain nor network-based analysis (i.e. resting-state networks relevant to social and motivational processing) of resting-state connectivity resulted in significant differences in similarity between friends and those farther removed in their social network. Likewise, exploratory analyses provided no evidence of neural homophily at rest. Specifically, graph theoretical measures of brain connectivity, including modularity, diversity and strength, were no more similar among friends than other more distantly connected pairs of students. Results from elastic-net regression, using the whole collection of local-level connectivity to predict social distance, also provide minimal evidence for similarity in resting-state functional connectivity among friends. Our findings were robust across individual cohorts of students and demonstrated a consistent non-significant result for homophily of resting-state connectivity. The inclusion of a data-driven approach to analysis (elastic-net regression) suggests our lack of evidence for the hypothesis is not due to noise from irrelevant variables or poor a priori selection of resting-state networks for dimensionality reduction. Likewise, results are not due to poor sampling from the overall population. Students participating in the fMRI component of the study exhibited demographic characteristics representative of their original class cohorts, from which 87% - 96% of enrolled students provided social network data.

Current literature supports a role for social closeness in synchronisation of neural activation. For example, students who report higher social closeness to one another who also engage in silent eye contact prior to class exhibit stronger pairwise brain-to-brain synchrony during class activities compared with those less close who engage in eye contact ^34^. This increased synchronisation between friends is evident even when friends are in the absence of one another; using functional MRI of individual students in a real-world social network, Parkinson and colleagues (2018) investigated synchronisation of neural activation during video clip viewing and found evidence for homophily at the neural response level. Brain regions where response similarity was associated with social network proximity included areas implicated in motivation, learning, affective processing, memory, attention, theory of mind and language processing ^8^, some behavioural traits of which have exhibited homophily in previous non-imaging studies ^4,5,35,36^. These results suggest that, at least in terms of cognitive processing, similarities in behaviour relate to similarities in brain function.

In contrast, the current study examined neural homophily during a resting state scan. Our findings suggest that neural homophily observed in previous work may be specific to stimulus-evoked activation, and may not extend to stimulus-free intrinsically-generated brain activities. Importantly, stimuli used by Parkinson and colleagues (2018) included video clips of comedy, debates and documentaries, intended to evoke social and emotional responses from participants. The homophily observed in their study may therefore be dependent on cognitive processes important for social interaction. This would also explain how our resting-state experiment, which was relatively devoid of social context, failed to elicit homophilic outcomes.

It should be noted that, in a stimulus-free environment, i.e. during rest, subjects are free to mind wander, providing no time-locked cue with which to directly compare activation between two subjects. Instead, simultaneous activation of spatially disparate brain regions is evaluated within each subject to identify networks of brain regions that exhibit highly correlated patterns of activity. Therefore, in the current study, we evaluated the correlational strength between every node in the brain for each individual participant and correlated the whole-brain or resting-state network connectivity pattern between every pair of participants in the social network. This was a powerful approach to evaluating dyadic similarities in resting-state brain function in a social network, but at the same time, this novel approach makes it difficult to directly compare the current findings with the previous one ^8^, which focused on the similarities of the activation pattern rather than the pattern of brain functional connectivity. Given the evidence that the architecture of task-based networks closely resembles networks seen at rest ^37^, it may be an interesting future inquiry to examine whether neural homophily is observed in the brain functional network connectivity triggered by external cues and stimuli.

A notable difference between our sample population and that of Parkinson et al. is the age at which social network and imaging data were collected (schoolchildren vs undergraduate students). As researchers have found associations between pubertal development and strength of intrinsic functional connectivity ^38^, our younger sample may exhibit less intrinsic network homophily than more mature samples due to greater brain variability between subjects.

In addition to intrinsic network strength, functional resting-state network architecture changes throughout the lifespan and can differ between adolescents and adults ^39^. For example, differences in DMN connectivity have been reported between adults and children ^40^ as well as within individuals throughout early adolescence ^41^. To ensure that the resting-state network maps used in our analyses were appropriate for the age group in our study, we ran independent components analysis in our student sample and compared our sample-derived independent components with the resting-state network maps used in our analyses. High levels of overlap were observed for salience, lFPN and rFPN maps and our sample-derived components. The DMN was represented over several components, though division of the DMN into smaller components in ICA is common.

The current study benefits from several strengths. Most notably, we evaluated homophily of resting-state connectivity in three different social networks comprising students of the same gender and similar age and education level, eliminating by design these demographic variables as possible sources of confound for neural homophily. Cohorts were evaluated as independent samples and then as individual studies in meta-analyses, ensuring sufficient statistical power of the overall analysis to find neural homophily. Homophily based on cognitive ability has been reported at a higher rate among girls compared with boys ^5^ and polygenic scores for educational achievement are more homogenous in women’s female social networks compared with men’s male social networks ^7^. As intelligence and cognitive control are reflected in the resting-state connectivity ^16^, we therefore anticipated that an all-female sample would exhibit stronger homophily of resting-state connectivity than a mixed-gender or all-male sample. Based on these attributes, our study was appropriately designed to identify homophily at the resting-state connectivity level.

These results contribute to the homophily literature by suggesting that homophily at the neural level may require some external stimulus that engages individuals in social or cognitive thoughts before synchronisation or similarities in connectivity are evident. To further our understanding of how our brain functioning is shaped by social factors, future research should examine the exact conditions under which neural homophily can be observed.

## Methods

### Participants

For the social network component of the study, individuals 12-14 years of age in years 8 (cohort 1: n=59; cohort 2: n=51) and 9 (cohort 3: n=65) were recruited from a private girls’ day and boarding school in the United Kingdom (as part of a larger study). Participants were recruited during 2017 and 2018 from total year group pools of 62 (cohort 1), 53 (cohort 2) and 75 (cohort 3) students, corresponding to inclusion rates of 95%, 96% and 87%, respectively.

The study was approved by the University Research Ethics Committee. All research was conducted in accordance with relevant guidelines, students gave informed written assent to take part in the study and consent was obtained from legal guardians.

Individuals from cohorts 1, 2 and 3 were also invited to take part in the functional magnetic resonance imaging (fMRI) component of the study. Twenty-eight students from cohort 1, 17 students from cohort 2 and 34 students from cohort 3 were recruited into the fMRI study (cohorts 1-fMRI, 2-fMRI and 3-fMRI, respectively); of these, 5 participants from cohort 1-fMRI and 6 from cohort 3-fMRI had unusable data due to artefacts caused by dental braces. Children with braces were excluded from cohort 2-fMRI at the time of screening. This resulted in the final inclusion of 68 students in the fMRI component of the study, consisting of 23 (cohort 1-fMRI, 12-13 years of age), 17 (2-fMRI, 12-13 years of age) and 28 (3-fMRI, 13-14 years of age) students in respective cohorts.

Exclusion criteria pertaining to all groups consisted of standard safety-related contraindications for MRI.

The study was well powered to detect an effect of social distance on neural similarity at rest, based on an in-house simulation (see Supplementary Materials: Study Power).

### Data acquisition

The social network data were acquired in class using an online survey administered with SurveyMonkey (SurveyMonkey Inc., San Mateo, California, USA). Social network data were acquired in October and November of 2017 (cohorts 1 and 3) and 2018 (cohort 2), one-to-two months after the start of the academic year. Students had been registered at the school for a maximum of one (cohorts 1 and 2) or two (cohort 3) years when the survey took place. Students were asked to rate how much time they voluntarily spent with each member of their year group (roster-and-rating method) as well as nominate up to five students in their year group whom they considered their “best friends”. Participants were asked to report on social interactions within their own year group only. As mentioned in the Results, nomination data gave similar outcomes to roster-and-rating and so is not discussed further. Investigators were blinded to the identity of students.

### Social network characterisation

Social network analysis was performed using the igraph package in R ^42,43^ (Details in Supplementary Materials). Data were included only for individuals who took part in the social network survey; any non-reciprocated ties or outgoing social ties with those not participating in the survey were removed.

### MRI analysis

Structural and resting-state functional MRI data were acquired using a Siemens Magnetom Prisma_fit 3T scanner. Details of MRI data acquisition and pre-processing are provided in the Supplementary Materials: Functional MRI Data Analysis.

Analysis of pre-processed data is illustrated in Figure 3. First, motion-corrected fMRI data were divided into 272 parcels using a whole-brain parcellation scheme ^44^ combining parcels from the Human Brainnetome Atlas ^45^ and the probabilistic MR atlas of the human cerebellum ^46^. Mean BOLD time series were extracted from each parcel and the Pearson’s correlation coefficient was determined for each pair of parcels (nodes) in MATLAB 2016a (MathWorks, USA). This produced a weighted, undirected whole-brain matrix of functional connectivity for each participant, which could be used for inter-subject correlations (all self-self nodal connections were removed and matrices were *z* transformed prior to further analysis). The *z*-transformed whole-brain matrix of subj_i_ was vectorised and correlated with the (vectorised) *z*-transformed whole-brain matrix of subj_j_, then subj_k_, then subj_l_, and so on. This method provides a measure of the similarity of connectivity strength in whole-brain networks within participant pairs. Prior to further analysis, correlation strengths were standardised within each cohort to have a mean of 0 and a standard deviation of 1.

Resting-state network comparisons were performed in a similar manner (see Supplementary Materials: Resting State Analysis and Supplementary Materials: Fig. S7).

### Main Analyses: Dyadic similarities in functional connectivity as a function of social proximity

To test our main hypotheses, we examined whether similarities in resting-state connectivity and brain network characterization were explained by social proximity. In this set of analyses, each student dyad (pair of students) in the fMRI cohort was described by two independent variables: social distance between the two students (e.g. distance of 1, 2 or 3) and whether or not their community affiliation was the same (‘yes’ (1) or ‘no’ (0)).

Using pairs of students (dyads) as the unit of analysis creates dependence in the data, caused by the involvement of every student in multiple dyads (cross-nesting) ^47 48^. Using ordinary least-square methods in such data potentially increases Type-1 error rates. We accounted for this dependence structure by including each dyad member (student) as a random variable in a linear mixed effects (LME) model with crossed random effects (analysed using the lme4 package ^49^ in R; see Supplementary Materials: LME Model Specification for more details and see ^50^, for a similar model specification). Any subject-specific effects on the dyadic outcomes are therefore accounted for in the model ^51^. Dependent variables were similarities in whole-brain connectivity and resting-state network (DMN, salience network, lFPN and rFPN) connectivity. For each dependent variable, we tested a model with social distance as the independent variable, and tested another model with community affiliation as the independent variable. Cohorts were initially analysed separately to preserve their independence (given that we did not measure social ties between the different year groups). We then integrated the regression coefficients of the LME models from three cohorts using a random-effect meta-analysis with the Metafor package ^52^ in R.

Effects of demographic data as well as their interactions with social distance or community affiliation were also included in a second series of models to determine whether boarding status (i.e. whether students matched in their boarding status [boarding or day student]) and ethnicity (i.e. whether or not students reported belonging to the same ethnic group) affected the relationship between social proximity and similarity in resting state connectivity between students. As only one student from each fMRI cohort was left-handed, we elected not to include handedness as a covariate.

### Exploratory analyses: graph theory measures of connectivity

For each individual in the fMRI cohorts, graph metrics of brain connectivity were derived using the Brain Connectivity Toolbox ^31^ in MATLAB (2016a). We were specifically interested in measures of brain modularity, node-level strength and node-level diversity, as these measures are designed to deal with the full interconnectedness of weighted functional brain networks ^53^. Each of these measures are defined further in the Supplementary Materials: Brain Network Characterisation.

As for the main analyses, graph measures of connectivity were compared between all student dyads within each cohort. Similarities in nodal strength and nodal diversity were determined by the Pearson’s correlation coefficient between two members of a dyad, whereas modularity (represented by a single value for each participant) was determined by the absolute difference between members in the dyad (see Supplementary Materials: **Error! Reference source not found.**S8 for details).

Methods for the data-driven approach to prediction of social proximity from neural similarity are provided in the Supplementary Materials.

## Acknowledgements

The authors would like to thank the students who gave their time to take part in the social network and imaging components of this study, as well as the parents, teachers and staff of Queen Anne’s School who made data collection possible. We also wish to thank the Centre for Integrative Neuroimaging and Neurodynamics at the University of Reading for assistance with scanning and timetabling. Many thanks also to the members of the Motivation Science Lab, University of Reading, who gave their time to assist with social network and imaging data collection.

## Additional Information

The authors declare they have no conflicts of interest.

## Data Availability

The datasets generated during and/or analysed during the current study are available from the corresponding author on reasonable request.

## Author contributions

C.B.M. Contributed to the design of the study and data collection, analysed data and wrote the manuscript; L.G.B. Contributed to the design of the study and collected social network and imaging data; A.F. contributed to the design of the study and data collection; N.M. contributed to the design of the study and data collection; L.F. contributed to data collection and statistical analysis; P.R. contributed to the design of the study; K.M. contributed to the design of the study and assisted with data collection and statistical analysis as well as interpretation of findings. All authors have read and approved the final manuscript.

