## Supplementary for "No evidence for a relationship between social closeness and similarity in resting-state functional brain connectivity in schoolchildren"

### Supplementary Methods

#### Study power

The fMRI sample affords a total of 767 dyads to examine potential neural homophily. Currently, there is no software available to perform a statistical power analysis with the mixed-effects model we specified to test our hypothesis with the dyadic data (see section on *Dyadic similarities in functional connectivity as a function of social proximity* for linear mixed effects model specification); as such, we created an in-house simulation code to perform a power analysis. The simulation results showed that our sample size is sufficient to detect very small effects of social distance on similarity. Specifically, the study was powered to detect a change in z standardised similarity score of at least .08 with one unit change in social distance at a power of 80%, and a change in *z*-standardized similarity score of .10 at a power of 95%. As previous neuroimaging work identified a difference of -.2 to -.23 in *z*-standardized similarity scores between social distances 1 to 2 and 2 to 3, respectively [1], we consider our sample size to be sufficiently powered to detect meaningful changes in similarity score between our social distance units.

**Social network characterisation**

Social network data were processed as follows: Roster-and-rating data (5-point scale) were binarised using a threshold of 4 (i.e. only instances in which students spent "more than some" or "most" of the time with another student were included). This threshold was selected to mitigate central tendency bias often reported with Likert-type questionnaires [2]. Any non-mutual connections were then removed (i.e. if $subj_i$ gave $subj_j$ a rating of 4 or greater but $subj_j$ gave $subj_i$ a rating below 4, the connection would be lost). This yielded an unweighted (binary), undirected (reciprocal) adjacency matrix for each cohort, from which social networks graphs were derived.

**Social Network Metrics**

Each student cohort was described in terms of its network characteristics, in particular, its network diameter, modularity, mean path length, reciprocity and density. Network diameter is the length of the longest geodesic distance between two nodes in the network, i.e. the number of edges between $subj_i$ and $subj_j$ when these individuals are the farthest from each other in the network.

Modularity ($Q$) is a measure of how easily a network segregates into smaller subnetworks and is defined by the equation:

$$Q = \frac{1}{2m}\sum_{ij}\left(A_{ij} - \frac{k_i * k_j}{2m}\right)\delta(c_i, c_j)$$

where $m$ is the number of edges in the network, $A_{ij}$ is the element of the $A$ adjacency matrix in row $i$ and column $j$, $k_i$ is the degree (number of edges associated with a node) of $i$, $c_i$ is the community to which $i$ belongs and $\delta(c_i, c_j)$ is 1 if $c_i = c_j$ and 0 otherwise. Nonzero values of $Q$ represent deviations from randomness; a value above 0.3 is an indicator of significant community structure in a network [3].

Mean path length is the mean shortest path length (number of edges separating a pair of nodes) between all nodes in the network. Reciprocity defines the proportion of connections in a directed graph that are mutual connections. It is otherwise defined as the probability that the counterpart (j to i) of a directed edge (i to j) is included in the graph. Graph density is the ratio of actual connections (edges) to possible connections in the graph; larger values denote more densely connected networks.

**Functional MRI Data Analysis**

All participants were imaged using a 32-channel head coil. A structural T1-weighted image was acquired using an MPRAGE sequence [4] with repetition time (TR) 2300 ms; echo time (TE) 2.29 ms; inversion time 900 ms; flip angle 8°; in-plane acceleration (GRAPPA) factor of 2; field of view (FOV) 240 mm; voxel size 0.9 x 0.9 x 0.9 mm.

Resting-state functional images were acquired over 10 minutes using the Siemens two-dimensional multiband gradient-echo echo planar image (EPI) sequence with TR 1500 ms, TE 30 ms; multiband slice acceleration factor 4; GRAPPA 2; flip angle 66°; echo spacing 0.93 ms; EPI factor 96; phase-encode direction posterior >> anterior; slices 68; volumes 400; FOV 192 mm; voxel size 2 x 2 x 2 mm.

During resting-state data acquisition, participants were asked to lie still with eyes open and look at a blank screen in front of the scanner. Instructions were given to relax and think of nothing in particular.

FMRI data processing was carried out using FEAT (FMRI Expert Analysis Tool, version 6.0 [5]) part of the FMRIB Software Library (FSL; Oxford, United Kingdom) [6,7]. Registration of functional images to high resolution structural and Montreal Neurological Institute (MNI-152) standard space images was carried out using FLIRT [8,9]. Registration from high resolution structural to standard space was then further refined using FNIRT nonlinear registration [10,11].

The following pre-statistics processing was applied: motion correction using MCFLIRT [9]; non-brain removal using BET [12]; multiplicative mean intensity normalization of the volume at each time point and high pass temporal filtering (Gaussian-weighted least-squares straight line fitting, with sigma=50.0s). Independent components analysis (ICA)-based exploratory data analysis was carried out using MELODIC [13], in order to investigate the possible presence of unexpected artefacts. FIX - FMRIB's ICA-based Xnoiseifier [14,15] was used to auto-classify ICA components into "good" and "bad" components following hand-classification and training using a sample of 10 subjects' data (4 from cohort 1- fMRI and 3 each from cohorts 2-fMRI and 3-fMRI, all randomly selected). "Bad" components were removed from the data and clean data were registered to standard space using warping parameters determined by FEAT. These data were then used for further analysis.

**Resting-state network analysis**

Default mode network (DMN) and frontoparietal networks (FPNs) were extracted from [16]. *Z*-statistic images of independent components analysis (ICA) maps were then binarised using a threshold of $z$=4 and warped to 1 mm MNI-152 standard space. Our threshold of $z$=4 is larger than that reported by [16], who adopted a threshold of $z$=3 for presentation of resting-state networks. Our larger $z$ threshold was chosen to increase the probability that brain nodes overlapping DMN and FPN spatial maps were truly part of these networks. The salience network was derived using the online meta-analysis tool Neurosynth.org [17] (accessed January 2019; seed positioned in the anterior insular [x=36, y=18, z=4]). The salience network was binarised using a threshold of 0.3, based on results from previous research [18]. The overlap between each resting-state network and the whole-brain parcellation was determined by applying a binarised mask of the resting-state network over whole-brain parcellation data. Nodes with at least 50 voxels overlapping the resting-state network were used in the analysis. A graphical depiction of the analysis is provided figure S7. Functional resting-state network architecture develops throughout the lifespan and can differ between adolescents and adults [19]. As we used resting-state network maps derived from mostly adult data, we wanted to ensure that these network maps were appropriate for use in our sample. We first ran independent components analysis (ICA) on resting-state fMRI data from our three student cohorts using MELODIC (Multivariate Exploratory Linear Optimized Decomposition into Independent Components) within the FMRIB Software Library (FSL). ICA maps were then compared to resting-state network maps derived from Neurosynth.org and Smith et al. 2009. All resting-state networks overlapped with independent components derived from our student sample. High levels of overlap were observed for salience, lFPN and rFPN maps, whereas the DMN was represented over several components. As all resting-state networks were identifiable in the study-derived ICA maps, they were deemed suitable for use in the analysis.

**LME model specification**

The LME model used in the current analysis is specified as:

$$Similarity_{ij} = \beta_{00} + \beta_{01} * social\ distance_{ij} + subj_i + subj_j + e_{ij}$$

$Similarity_{ij}$ is the similarity in brain function between students $i$ and $j$, where $\beta_{00}$ is the intercept, and $\beta_{01}$ represents the relationship between social distance and brain similarity; $subj_i$ and $subj_j$ are crossed random effects (i.e. student-specific effects) of students $i$ and $j$, respectively, and $e_{ij}$ are residuals. We posited a Gaussian distribution for the random effects and residuals. No distribution assumption was made for social distance. Where social network data were described using community affiliation, $social\ distance_{ij}$ in the above equation was replaced with $community\ similarity_{ij}$ (i.e. whether or not two students belong to the same friendship community, as determined by the Louvain method).

We also tested a model with random slopes (i.e. random slopes of the subjects were added to the model above). This model failed to converge in most of the analyses. The omission of random slopes could make the statistical test less conservative [20,21]. However, as our results all showed non-significant effects in the model without random slopes, we decided not to pursue the model with random slopes any further.

**Brain network characterisation**

For graph theory analysis of functional connectivity, modularity was defined as:

$$Q^* = Q^+ + \frac{v^-}{v^+ + v^-} Q^-$$

$$= \frac{1}{v^+} \sum_{ij} \left(w_{ij}^+ - e_{ij}^+\right) \delta\left(M_i, M_j\right) - \frac{1}{v^+ + v^-} \sum_{ij} \left(w_{ij}^- - e_{ij}^-\right) \delta\left(M_i, M_j\right)$$

where $Q^+$ is modularity from positively weighted connections, $Q^-$ is modularity from negatively weighted connections, $v^+$ is the sum of all positive connection weights of $i$, $v^-$ is the sum of all negative connection weights of $i$, $w_{ij}^+$ is the present within-module positive connection weights, $w_{ij}^-$ is the present within-module negative connection weights, $e_{ij}^+$ is the chance expected within-module positive connection weights, $e_{ij}^-$ is the chance expected within-module negative connection weights and $\delta\left(M_i, M_j\right)$ is 1 if $i$ and $j$ are in the same module and 0 otherwise.

The absolute difference in brain modularity within a dyad pair (i.e. between $subj_i$ and $subj_j$) was calculated for every pair of dyads in a social network. Differences were then standardised within each cohort to have a mean of 0 and standard deviation of 1.

Nodal strength (a node-level measure of centrality, the importance of a node in its network) and diversity (a node-level measure of integration that takes into account the strength of a node within its own module) were also calculated using asymmetric values for positively and negatively weighted connections, as previously described by Rubinov & Sporns (2011). Strength was defined as:

$$S_i'^{*} = s_i'^{+} - (\frac{s_i^-}{s_i^+ + s_i^-})s_i'^{-}$$

where $s_i'^{+}$ is the normalised sum of positive connection weights associated with node $i$, and $s_i^+$ and $s_i^-$ are the raw sums of positive and negative connections weights, respectively, associated with node $i$.

Diversity was defined as:

$$h_i^{*} = h_i^{+} - (\frac{s_i^-}{s_i^+ + s_i^-})h_i^{-}$$

and

$$h_i^{\pm} = \frac{1}{\log m}\sum_{u \in M} p_i^{\pm}(u) \log p_i^{\pm}(u)$$

where $p_i^{\pm}(u) = \frac{s_i^{\pm}(u)}{s_i^{\pm}}$, $s_i^{\pm}(u)$ is the strength of node $i$ within module $u$ (the total weight of connections of $i$ to all nodes in $u$) and $m$ is the number of modules in modularity partition $M$.

Nodal strength and diversity data from each participant were compared with every other participant from the fMRI cohort using a Pearson's correlation. Prior to further analysis, correlation strengths were standardised within each cohort to have a mean of 0 and a standard deviation of 1.

**Data-driven predictive model of social proximity from neural similarity**

To examine whether brain functional connectivity encodes social distance, we employed regularised regression techniques to predict social distance between two students based on similarities in their functional brain connectivity of all pairs of nodes. Specifically, we computed the absolute difference in connection strength for each edge in the 272 node brain network (i.e. each node-to-node connection) for every student dyad in the fMRI cohort. This yielded $Nnodes * (Nnodes - 1)/2 = 36{,}856$ similarity (absolute difference) measures for each student dyad, where $Nnodes$ is the number of nodes in the whole-brain parcellation (i.e. 272). We then employed elastic-net regularised linear regression to predict social distance of student dyads from the 36,856 similarity measures of resting-state functional connectivity.

Elastic-net regression combines penalty features of lasso and ridge regression techniques to shrink the coefficients of some regressors toward zero to deal with high dimensional data. Whereas lasso regression can shrink unnecessary regressors to zero and thereby reduce the number of predictors, ridge regression retains all regressors for inclusion in the model. Both lasso and ridge regression techniques have been shown to perform well when dealing with high-dimensional data under various conditions [22]. Combining the two approaches, elastic-net regression allows for adjustment of the lasso-to-ridge ratio (α), providing greater opportunity for better model fits [23].

Elastic-net regressions of connectivity and social distance data were conducted in R using the glmnet package [24]. Three regression models were trained using data from two fMRI cohorts each (see Table $S1$). One fMRI cohort's data were withheld during training so that the performance of the regression model could be evaluated using a previously unseen set of data.

Table S1. Training and testing sets for each predictive model. Each model was trained using data from two of the MRI cohorts and trained on unseen data from the third MRI cohort. Tuning parameters were determined separately for each model during the training phase.

| | Model 1 | Model 2 | Model 3 |
|---|---|---|---|

| | | | |
|---|---|---|---|
| Training set | Cohort 2-fMRI<br>Cohort 3-fMRI | Cohort 1-fMRI<br>Cohort 3-fMRI | Cohort 1-fMRI<br>Cohort 2-fMRI |
| Testing set | Cohort 1-fMRI | Cohort 2-fMRI | Cohort 3-fMRI |
| Tuning Parameters: | | | |
| λ | 209.085 | 176.565 | .068 |
| α | 0 | 0 | 1 |

The best-performing regression model for each training set was determined by optimising the tuning parameters λ and α (see Table 1). λ is a tuning parameter for the shrinkage penalty used to adjust the regression coefficients in the elastic-net regression. When λ = 0, the penalty term has no effect but as λ tends toward infinity, the shrinkage penalty grows, and the regression coefficient estimates approach zero. The optimal value of λ for each regression model was determined using a 10-fold nested cross-validation within the training data. The largest value of λ such that the cross validation error was within one standard error of the minimum was selected and the model was re-fit using all available observations.

The tuning parameter α dictates the ratio of ridge to lasso in the elastic-net regression. α values between 0 and 1 (with iterations of 0.1) were evaluated for each regression model. The best performing α value was chosen based on the smallest root mean squared error (RMSE) between predicted and observed values of social distance in the training data. Best-performing α values were α = 0 (pure ridge regression) for models 1 & 2 and α = 1 (pure lasso regression) for model 3. The performance of the optimised regression model was then evaluated by predicting the social distance of dyads in the previously unseen testing dataset.

After obtaining the predicted social distance scores from the elastic-net regression models, we evaluated the accuracy of predictive models by examining the relationship between the predicted social distance and the observed (actual) social distance. As before, there is dependency in the data structure, owing to the involvement of each student in multiple dyads, potentially inflating the test statistics assessing statistical significance of the accuracy scores. To account for the dyadic nature of the data, we again used an LME model to obtain *p* value using the observed social distance as the dependent variable and the predicted social distance as the independent variable, with $subj_i$ and $subj_j$ included as crossed random effects.

Finally, to evaluate the overall predictability of social distance using similarity in resting-state connectivity, we conducted a meta-analysis of LME models from the three elastic-net regressions (models 1, 2 and 3). A positive beta weight with 95% confidence interval excluding zero would indicate that prediction of social distance from neural similarity is feasible.

**Supplementary Results: Prediction of social distance based on node-to-node neural similarities**

Using elastic-net regression, we sought to determine whether similarities in node-to-node connectivity within the brain could predict social distance between students. Data were split into training and testing sets such that each cohort was used to train two models and test the predictive validity of a third model. Assignments of training and testing sets for the three models are provided in Table 1. Optimal parameters (i.e. α and λ values) for each model were determined using training data; these parameters were then used to predict social distance in the testing set (to which the model was naïve).

RMSE quantifies how much a set of predicted values differ from their observed counterparts by measuring the standard deviations of the prediction errors. Lower values indicate smaller errors. RMSEs were 0.60, 0.65 and 0.82 for models 1, 2 and 3, respectively. Correlations for the observed vs predicted distance with the *p*-values of the beta weights obtained from the LMEs between each pair of participants are presented in Figure S8a. Meta-analysis of LME models showed poor predictive power of models to classify social distance of dyads based on whole-brain functional connectivity

(Figure S8b). Our meta-analysis results suggest that our predictive models will not extrapolate well to predict social distance in previously unseen social networks.

### Supplementary Results: Tables and Figures

*Table S2. Fixed effects t values and corresponding p values for beta weights (slopes) from linear mixed effects models with crossed random effects for similarity in brain function based on social distance and community affiliation. Degrees of freedom were rounded to the nearest whole number.*

| Social network characteristic (fixed effects) | Functional connectivity metric | Year group | Degrees of freedom (df) | *t* | *p* |
|---|---|---|---|---|---|
| Social distance | Whole-brain network | Cohort 1 | 213 | -1.58 | 0.115 |
| | | Cohort 2 | 107 | 1.76 | 0.082 |
| | | Cohort 3 | 337 | -1.19 | 0.234 |
| | Default mode network | Cohort 1 | 229 | 0.68 | 0.495 |
| | | Cohort 2 | 109 | 1.01 | 0.317 |
| | | Cohort 3 | 343 | -1.73 | 0.085 |
| | Salience network | Cohort 1 | 217 | 0.73 | 0.467 |
| | | Cohort 2* | 110 | 0.52 | 0.606 |
| | | Cohort 3 | 338 | -1.51 | 0.131 |
| | Left frontoparietal network | Cohort 1 | 217 | 0.71 | 0.476 |
| | | Cohort 2 | 110 | 0.47 | 0.641 |
| | | Cohort 3 | 338 | -1.06 | 0.290 |
| | Right frontoparietal network | Cohort 1 | 219 | 0.63 | 0.532 |
| | | Cohort 2 | 110 | 0.61 | 0.544 |
| | | Cohort 3 | 339 | -1.03 | 0.305 |
| | Nodal strength | Cohort 1 | 220 | -0.23 | 0.820 |
| | | Cohort 2 | 112 | 0.42 | 0.672 |
| | | Cohort 3 | 347 | -0.61 | 0.539 |
| | Nodal diversity | Cohort 1* | 235 | -0.73 | 0.463 |
| | | Cohort 2 | 120 | 1.33 | 0.185 |
| | | Cohort 3 | 358 | 0.60 | 0.546 |

| | | | | | |
|---|---|---|---|---|---|
| | Brain modularity | Cohort 1 | 235 | -0.56 | 0.579 |
| | | Cohort 2 | 117 | -0.90 | 0.372 |
| | | Cohort 3 | 349 | 0.66 | 0.509 |
| Community affiliation | Whole-brain network | Cohort 1 | 214 | -0.03 | 0.972 |
| | | Cohort 2 | 105 | -0.50 | 0.621 |
| | | Cohort 3 | 333 | 1.29 | 0.198 |
| | Default mode network | Cohort 1 | 232 | -0.67 | 0.504 |
| | | Cohort 2 | 106 | -0.71 | 0.481 |
| | | Cohort 3 | 336 | 0.57 | 0.568 |
| | Salience network | Cohort 1 | 220 | -0.81 | 0.421 |
| | | Cohort 2 | 106 | -0.92 | 0.361 |
| | | Cohort 3 | 334 | 0.08 | 0.939 |
| | Left frontoparietal network | Cohort 1 | 220 | -0.70 | 0.485 |
| | | Cohort 2 | 106 | -0.60 | 0.551 |
| | | Cohort 3 | 333 | 0.32 | 0.751 |
| | Right frontoparietal network | Cohort 1 | 221 | -0.71 | 0.477 |
| | | Cohort 2 | 106 | -0.54 | 0.590 |
| | | Cohort 3 | 334 | 0.40 | 0.691 |
| | Nodal strength | Cohort 1 | 222 | -0.28 | 0.776 |
| | | Cohort 2 | 107 | 0.41 | 0.686 |
| | | Cohort 3 | 338 | 0.31 | 0.757 |
| | Nodal diversity | Cohort 1 | 237 | -1.07 | 0.285 |
| | | Cohort 2 | 110 | -0.37 | 0.709 |
| | | Cohort 3 | 343 | 1.19 | 0.235 |
| | Brain modularity | Cohort 1 | 238 | 0.92 | 0.357 |
| | | Cohort 2 | 102 | -0.23 | 0.818 |
| | | Cohort 3 | 335 | -0.72 | 0.470 |

*Model failed to converge

*Table S3. Fixed effects t values and corresponding p values for beta weights (slopes) from linear mixed effects models with crossed random effects. Results are presented for boarding status and ethnicity as well as interaction terms of boarding status and ethnicity with social closeness; df = degrees of freedom.*

| Social network characteristic (fixed effects) | Functional connectivity metric | Year group | Boarding status | | | Social closeness*boarding interaction | | | Ethnicity | | | Social closeness *ethnicity interaction | | |
|---|---|---|---|---|---|---|---|---|---|---|---|---|---|---|
| | | | *df* | *t* value | *p* value | *df* | *t* | *p* | *df* | *t* value | *p* value | *df* | *t* | *p* |
| Social distance | Whole-brain network | Cohort 1 | 212 | -0.60 | 0.551 | 209 | 0.56 | 0.576 | 203 | 2.63 | 0.009 | 213 | -0.17 | 0.865 |
| | | Cohort 2 | 104 | 2.83 | 0.006 | 105 | -3.07 | 0.003 | 109 | 0.32 | 0.750 | 106 | -0.03 | 0.975 |
| | | Cohort 3 | 335 | 0.18 | 0.859 | 337 | 0.07 | 0.947 | 343 | -1.06 | 0.288 | 331 | 1.81 | 0.072 |
| | Default mode network | Cohort 1 | 230 | -1.96 | 0.051 | 224 | 1.57 | 0.118 | 164 | 2.18 | 0.031 | 219 | -0.56 | 0.577 |
| | | Cohort 2 | 105 | 0.38 | 0.708 | 107 | -0.35 | 0.729 | 113 | -0.80 | 0.423 | 109 | 0.37 | 0.712 |
| | | Cohort 3 | 341 | 0.15 | 0.878 | 343 | 0.31 | 0.758 | 350 | 0.04 | 0.971 | 335 | 0.27 | 0.789 |
| | Salience network | Cohort 1 | 219 | -1.24 | 0.215 | 211 | 1.19 | 0.237 | 138 | 1.80 | 0.074 | 208 | -1.11 | 0.268 |
| | | Cohort 2 | 106 | 0.33 | 0.745 | 108 | -0.51 | 0.609 | 113 | -0.92 | 0.361 | 110 | 0.43 | 0.671 |
| | | Cohort 3 | 337 | -0.52 | 0.601 | 339 | 1.02 | 0.310 | 344 | 0.21 | 0.832 | 333 | 0.12 | 0.906 |
| | Left frontoparietal network | Cohort 1 | 218 | -1.27 | 0.207 | 211 | 1.31 | 0.192 | 145 | 1.77 | 0.079 | 209 | -1.24 | 0.217 |
| | | Cohort 2 | 106 | 0.29 | 0.770 | 108 | -0.33 | 0.742 | 113 | -1.11 | 0.270 | 109 | 0.71 | 0.479 |
| | | Cohort 3 | 336 | -0.65 | 0.516 | 338 | 1.25 | 0.212 | 344 | -0.08 | 0.935 | 332 | 0.41 | 0.685 |
| | Right frontoparietal network | Cohort 1 | 220 | -1.12 | 0.263 | 213 | 1.20 | 0.233 | 145 | 1.52 | 0.131 | 211 | -0.73 | 0.468 |
| | | Cohort 2 | 106 | 0.24 | 0.812 | 108 | -0.33 | 0.739 | 114 | -1.19 | 0.238 | 110 | 0.83 | 0.406 |

| | | | | | | | | | | | | | | |
|---|---|---|---|---|---|---|---|---|---|---|---|---|---|---|
| | | Cohort 3 | 337 | -0.34 | 0.738 | 339 | 0.86 | 0.389 | 345 | -0.17 | 0.867 | 333 | 0.64 | 0.524 |
| | Nodal strength | Cohort 1 | 219 | -0.67 | 0.502 | 214 | 0.52 | 0.605 | 183 | 3.51 | 0.001 | 215 | -1.49 | 0.138 |
| | | Cohort 2 | 107 | 2.92 | 0.004 | 109 | -3.02 | 0.003 | 118 | 0.38 | 0.708 | 112 | -0.04 | 0.971 |
| | | Cohort 3 | 344 | -0.98 | 0.327 | 347 | 0.72 | 0.472 | 355 | -0.84 | 0.400 | 336 | 0.92 | 0.357 |
| | Nodal diversity | Cohort 1 | 235 | -1.54 | 0.126 | 229 | 1.20 | 0.230 | 194 | 3.24 | 0.001 | 231 | -0.54 | 0.590 |
| | | Cohort 2 | 113 | 0.95 | 0.342 | 116 | -0.93 | 0.357 | 127 | -1.62 | 0.108 | 122 | 1.62 | 0.107 |
| | | Cohort 3 | 351 | -0.64 | 0.525 | 355 | 0.51 | 0.608 | 363 | -0.70 | 0.483 | 338 | 1.19 | 0.233 |
| | Brain modularity | Cohort 1 | 234 | 0.88 | 0.380 | 227 | -0.84 | 0.402 | 216 | 1.23 | 0.220 | 224 | -0.99 | 0.323 |
| | | Cohort 2 | 106 | -2.10 | 0.038 | 112 | 2.26 | 0.026 | 129 | 0.27 | 0.791 | 119 | 0.33 | 0.738 |
| | | Cohort 3 | 345 | 0.56 | 0.576 | 349 | -0.15 | 0.880 | 356 | 0.88 | 0.381 | 333 | -0.67 | 0.501 |
| Community affiliation | Whole-brain network | Cohort 1* | 225 | 0.11 | 0.910 | 209 | -0.23 | 0.821 | 119 | 2.52 | 0.013 | 211 | 1.65 | 0.101 |
| | | Cohort 2 | 106 | -1.28 | 0.205 | 106 | 1.52 | 0.132 | 121 | 0.65 | 0.518 | 105 | -1.05 | 0.297 |
| | | Cohort 3 | 333 | 1.06 | 0.290 | 334 | -0.56 | 0.573 | 371 | 1.34 | 0.181 | 328 | -1.20 | 0.231 |
| | Default mode network | Cohort 1 | 220 | -0.81 | 0.419 | 228 | -1.20 | 0.231 | 21 | 2.95 | 0.008 | 233 | 0.56 | 0.579 |
| | | Cohort 2 | 109 | 0.11 | 0.916 | 109 | -0.05 | 0.961 | 129 | -0.87 | 0.383 | 107 | -0.43 | 0.665 |
| | | Cohort 3 | 337 | 1.86 | 0.064 | 340 | -0.85 | 0.395 | 371 | 0.12 | 0.908 | 331 | 1.09 | 0.278 |
| | Salience network | Cohort 1 | 243 | -0.13 | 0.897 | 214 | -0.63 | 0.529 | 36 | 1.12 | 0.272 | 210 | 1.20 | 0.230 |
| | | Cohort 2 | 110 | -0.12 | 0.905 | 110 | -0.29 | 0.776 | 129 | -0.99 | 0.324 | 107 | -0.34 | 0.733 |
| | | Cohort 3 | 334 | 1.75 | 0.080 | 336 | -0.73 | 0.466 | 371 | 0.08 | 0.933 | 330 | 1.37 | 0.172 |
| | Left frontoparietal network | Cohort 1 | 243 | 0.01 | 0.994 | 213 | -0.54 | 0.591 | 42 | 0.91 | 0.369 | 210 | 1.42 | 0.156 |

| | | | | | | | | | | | | | |
|---|---|---|---|---|---|---|---|---|---|---|---|---|---|
| | Cohort 2 | 110 | 0.21 | 0.831 | 110 | -0.38 | 0.705 | 129 | -0.88 | 0.382 | 107 | -0.36 | 0.723 |
| | Cohort 3* | 334 | 2.13 | 0.034 | 335 | -1.20 | 0.229 | 370 | -0.09 | 0.924 | 329 | 1.50 | 0.134 |
| Right frontoparietal network | Cohort 1 | 243 | 0.04 | 0.970 | 215 | -0.24 | 0.812 | 42 | 1.06 | 0.294 | 213 | 1.25 | 0.214 |
| | Cohort 2 | 110 | -0.15 | 0.882 | 110 | -0.08 | 0.938 | 130 | -0.85 | 0.394 | 108 | -0.29 | 0.771 |
| | Cohort 3 | 335 | 1.83 | 0.069 | 336 | -0.91 | 0.363 | 371 | 0.21 | 0.836 | 330 | 1.21 | 0.229 |
| Nodal strength | Cohort 1 | 244 | -0.08 | 0.937 | 216 | -0.97 | 0.334 | 64 | 3.16 | 0.002 | 215 | 0.68 | 0.496 |
| | Cohort 2 | 114 | -1.13 | 0.261 | 114 | 2.01 | 0.046 | 131 | 0.88 | 0.381 | 109 | -0.85 | 0.398 |
| | Cohort 3 | 340 | 0.58 | 0.561 | 342 | -2.01 | 0.045 | 366 | 0.99 | 0.324 | 332 | -2.49 | 0.013 |
| Nodal diversity | Cohort 1 | 225 | -0.71 | 0.481 | 231 | 0.14 | 0.890 | 27 | 5.43 | 0.000 | 238 | 0.48 | 0.628 |
| | Cohort 2 | 124 | -0.60 | 0.553 | 124 | 1.12 | 0.267 | 100 | -0.46 | 0.644 | 117 | 0.04 | 0.965 |
| | Cohort 3 | 346 | 0.21 | 0.836 | 347 | -1.34 | 0.180 | 330 | 1.56 | 0.120 | 331 | -2.28 | 0.023 |
| Brain modularity | Cohort 1 | 228 | 0.38 | 0.708 | 230 | -0.87 | 0.388 | 60 | 0.79 | 0.430 | 227 | -0.16 | 0.874 |
| | Cohort 2 | 126 | 1.76 | 0.082 | 124 | -2.67 | 0.009 | 94 | 1.38 | 0.172 | 114 | -0.27 | 0.787 |
| | Cohort 3 | 340 | 0.44 | 0.660 | 345 | 0.73 | 0.463 | 321 | 0.12 | 0.906 | 329 | 0.91 | 0.363 |

*Model failed to converge for boarding model

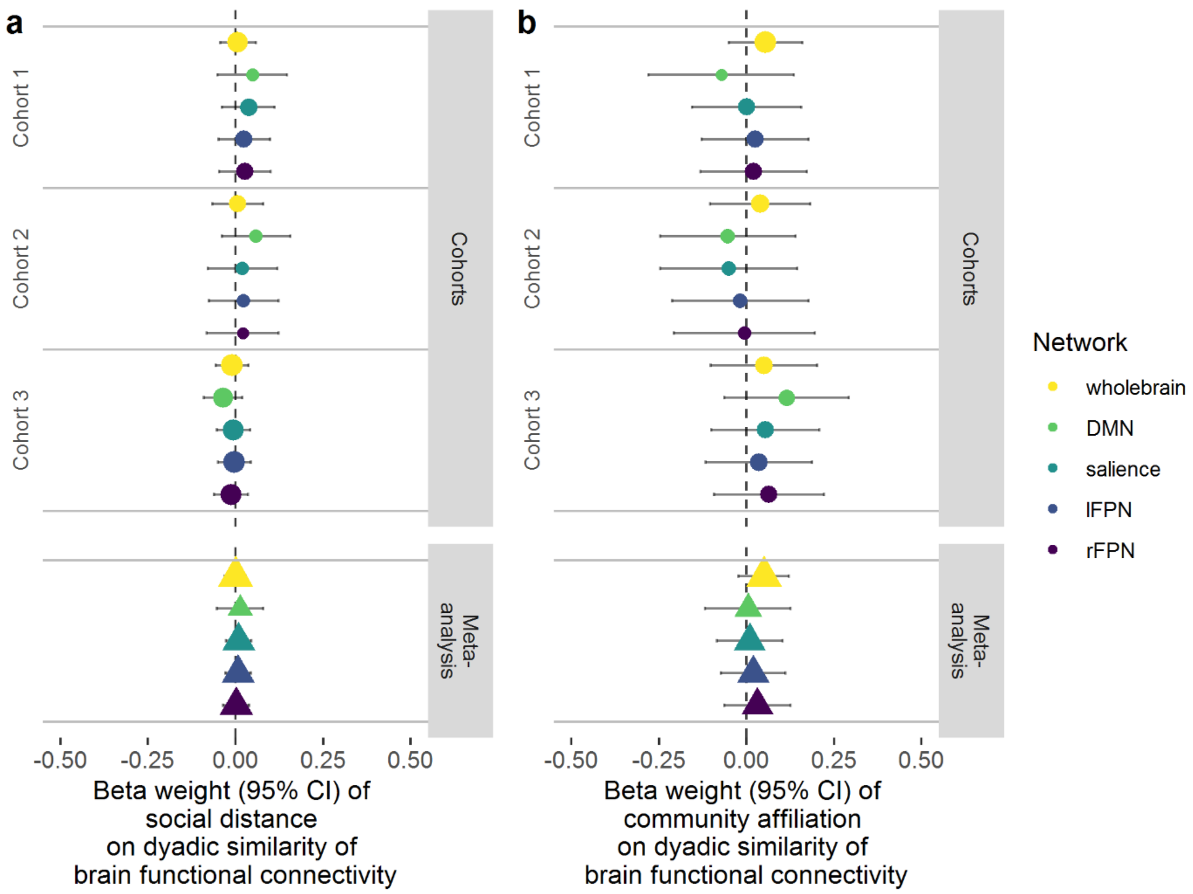


*Figure S1. Meta-analysis of brain similarity as a function of social distance (a) and community affiliation (b) for nominated friendships*

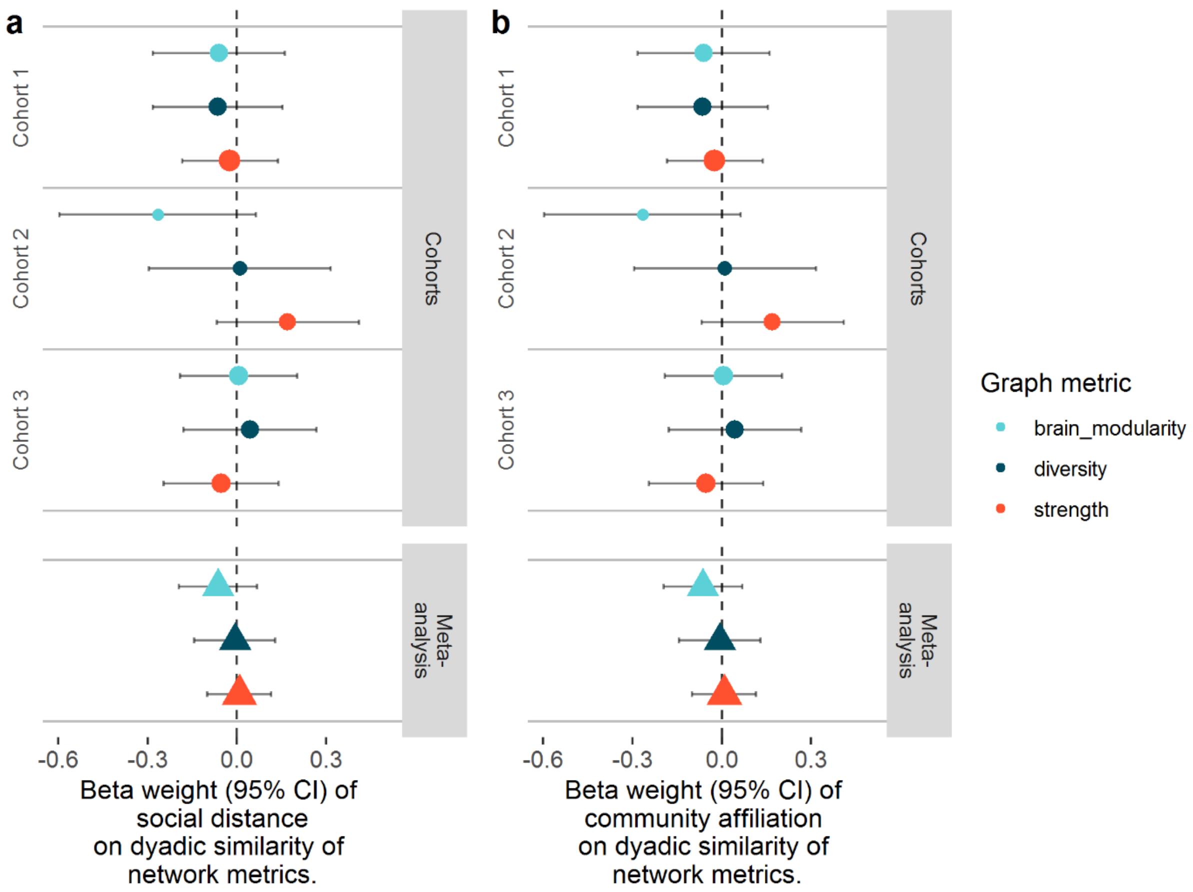


*Figure S2. Meta-analysis of graph metric similarity as a function of social distance (a) and community affiliation (b) for nominated friendships*

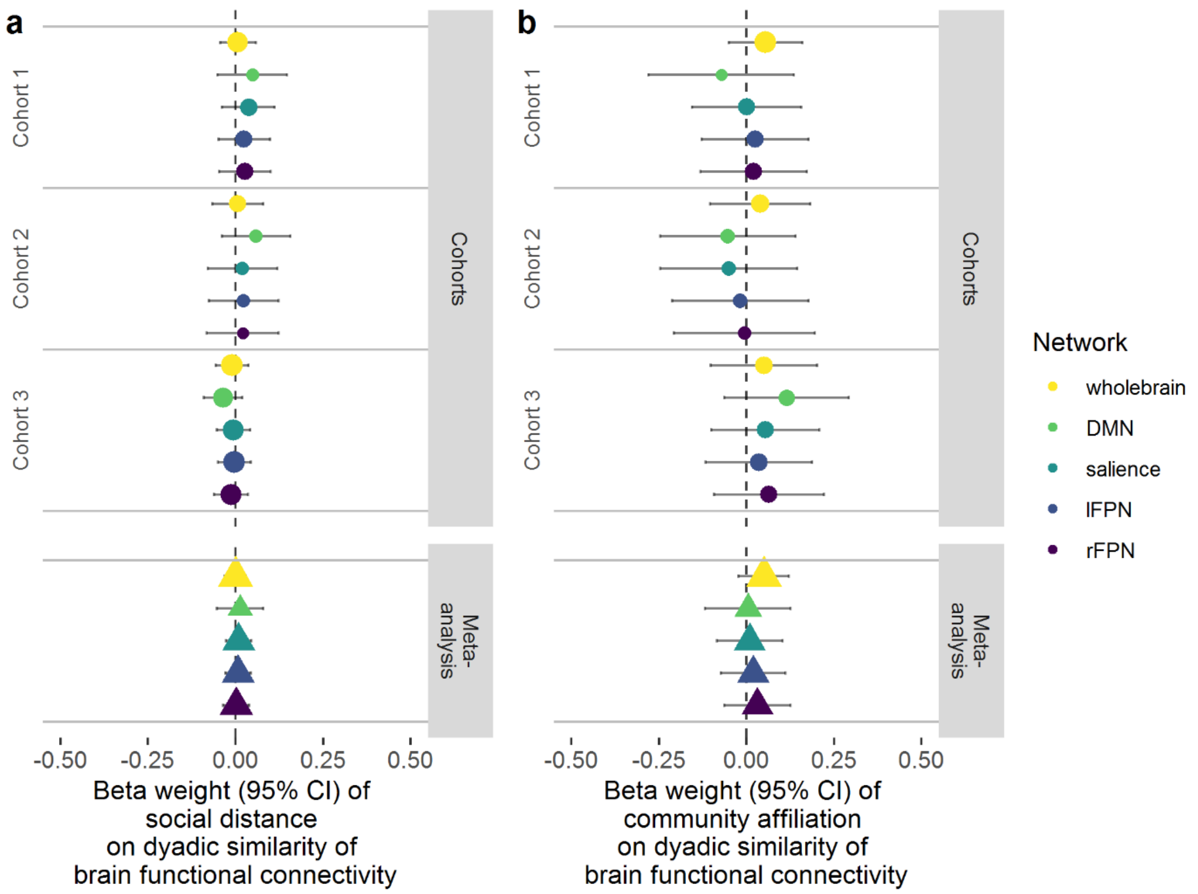


*Figure S3. Meta-analysis of brain similarity as a function of social distance (a) and community affiliation (b) for roster-and-rating method, threshold 5 (I spend "most of my time" with this person)*

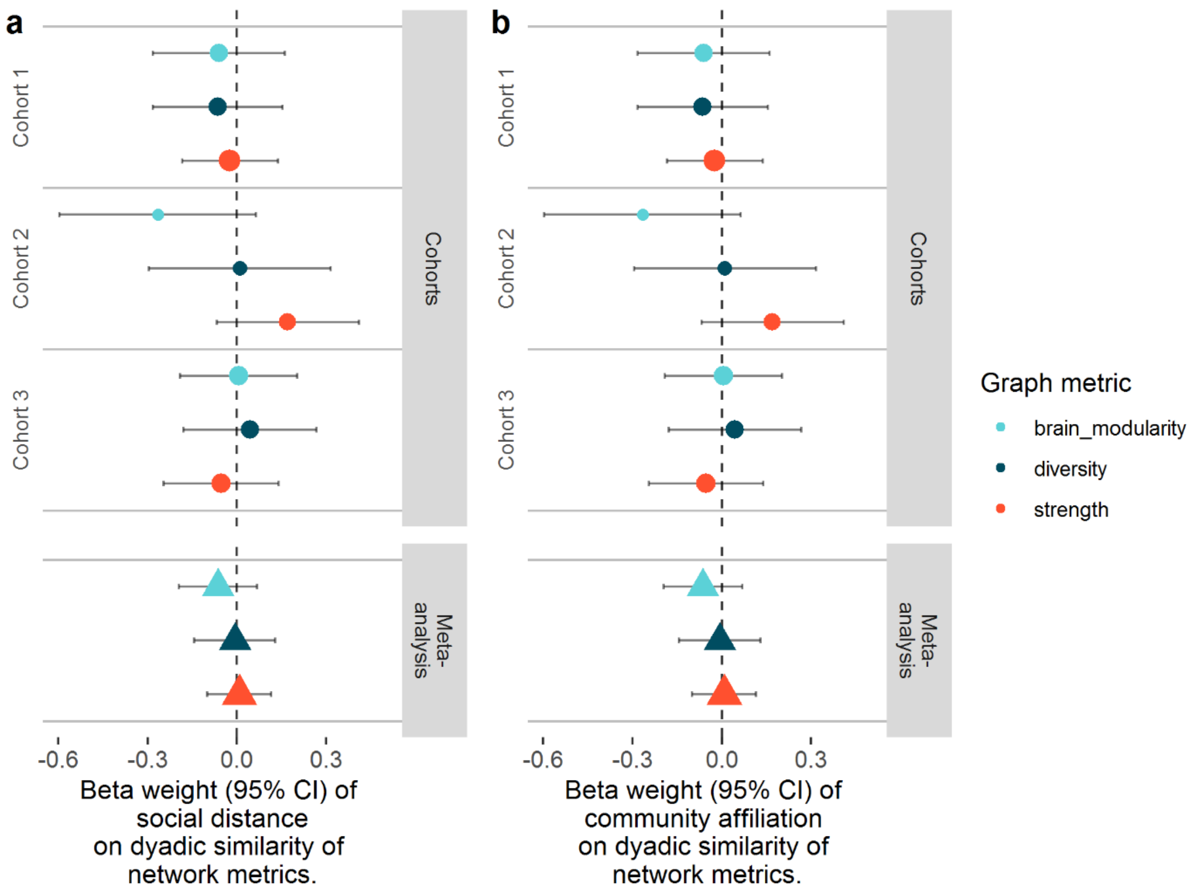


*Figure S4. Meta-analysis of graph metric similarity as a function of social distance (a) and community affiliation (b) for roster-and-rating method, threshold 5 (I spend "most of my time" with this person).*

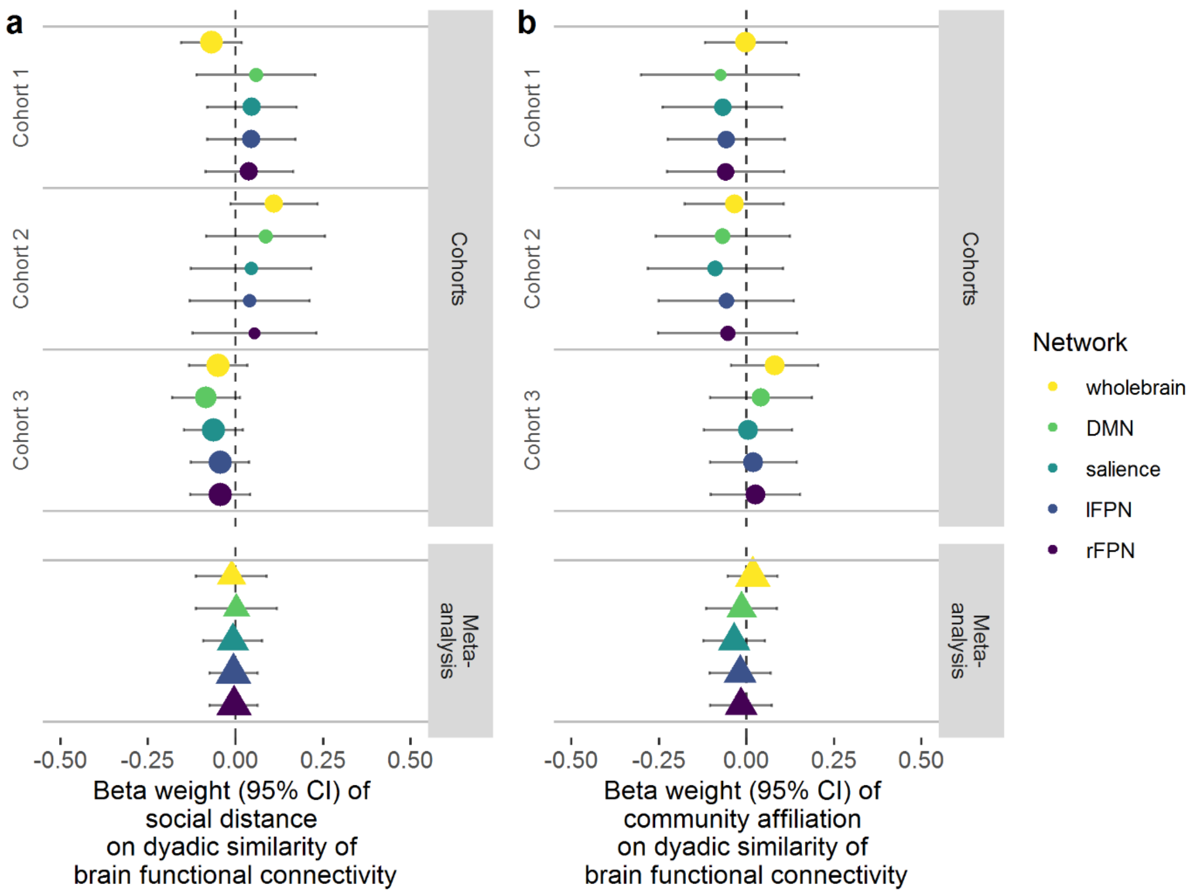


*Figure S5. Meta-analysis of brain similarity as a function of social distance (a) and community affiliation (b) for directed (non-mutual) friendship networks.*

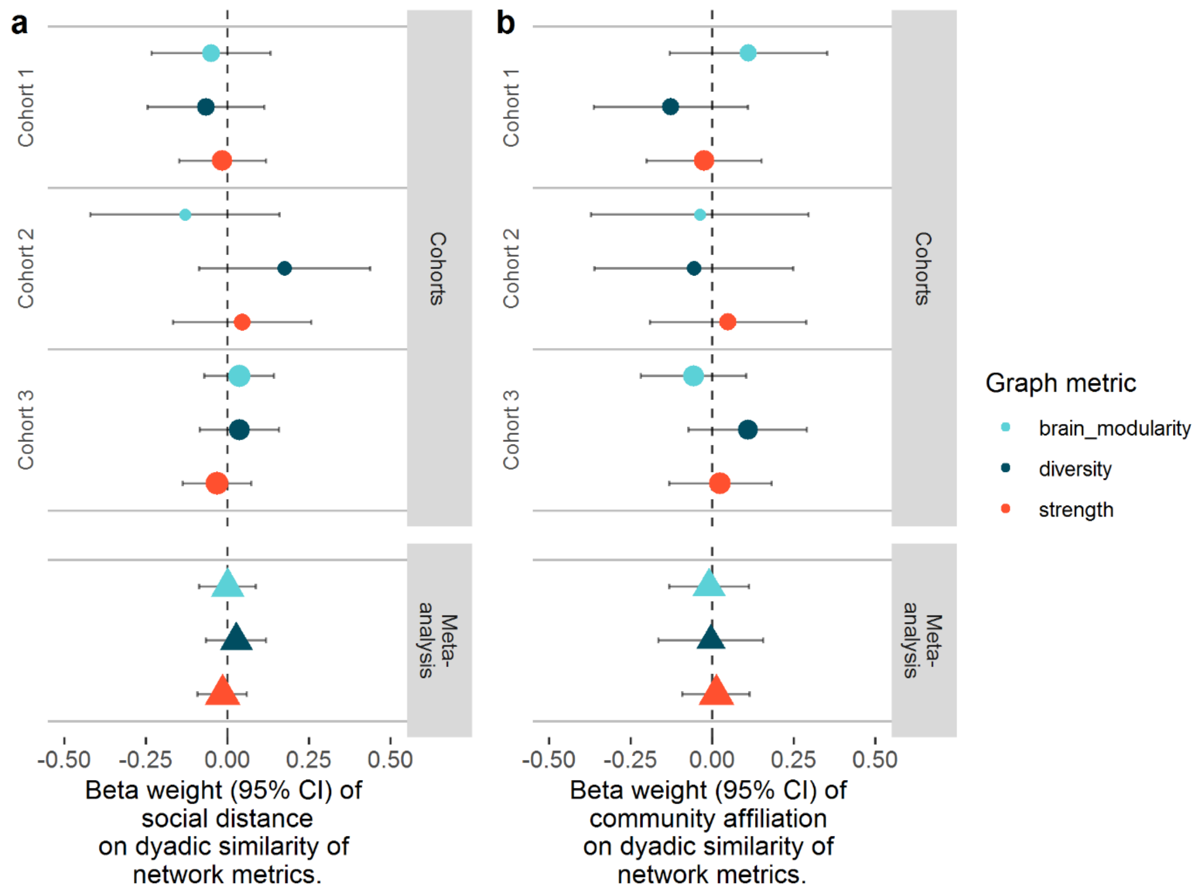


*Figure S6. Meta-analysis of graph metrics as a function of social distance (a) and community affiliation (b) for directed (non-mutual) friendship networks.*

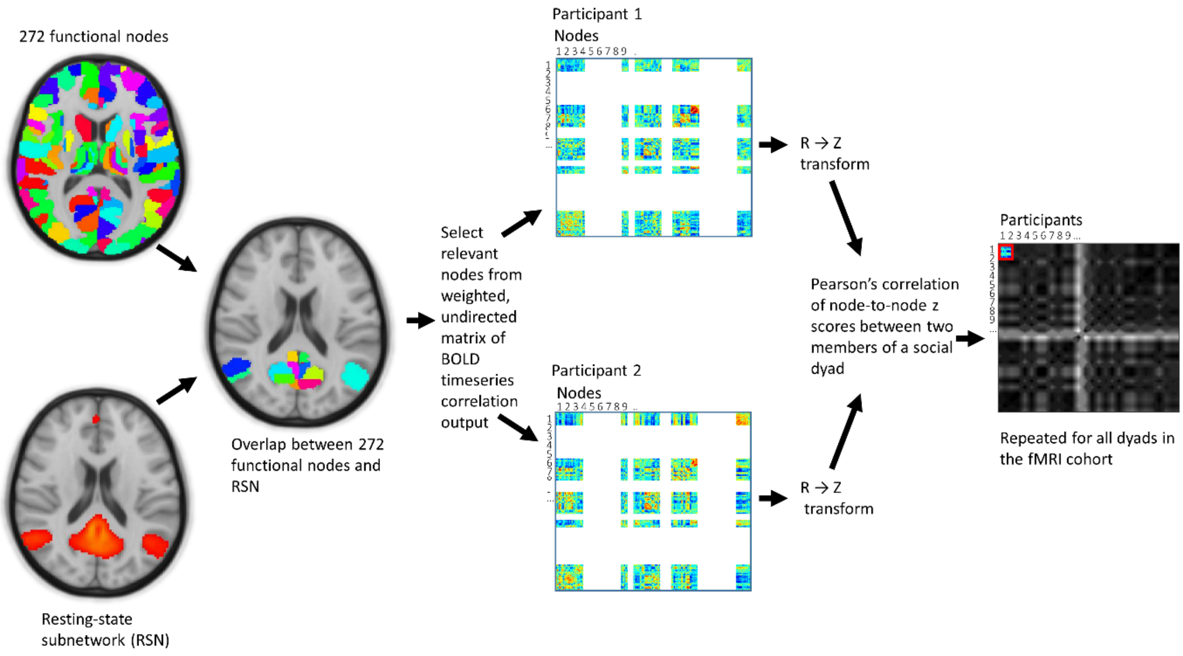


*Figure S7. Processing pipeline for determining similarity of functional connectivity between participants' resting-state networks.*

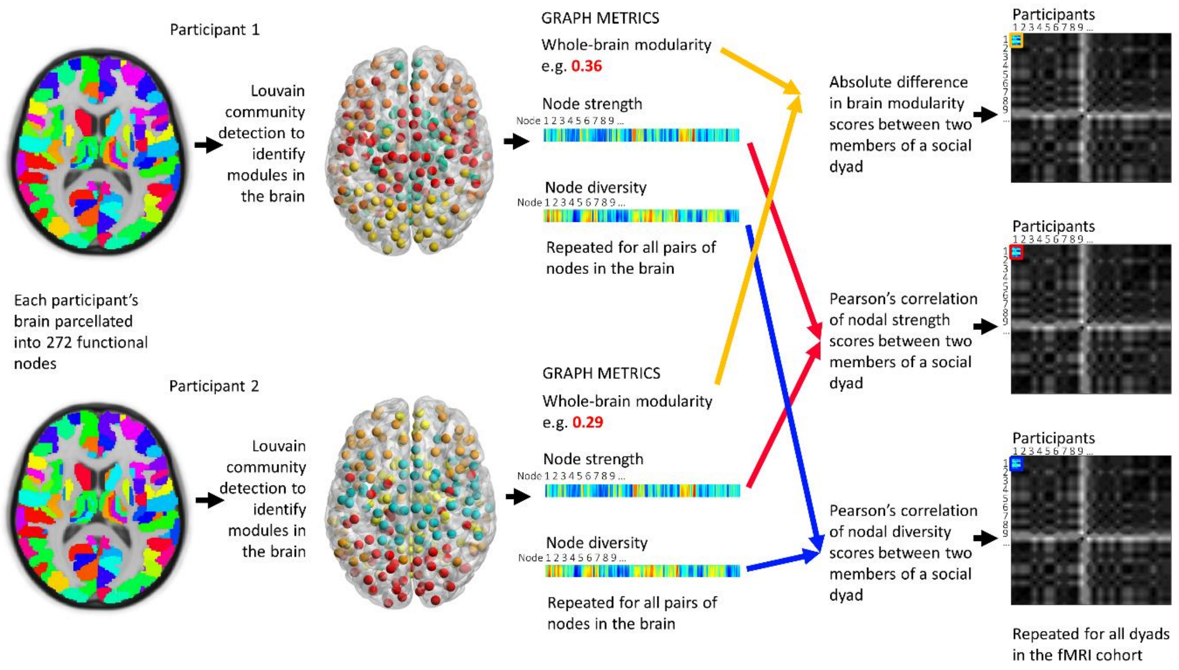


*Figure S8. Processing pipeline for graph metric analysis of resting-state data. Community detection was performed on each participant's brain data to determine modules (communities) from which further graph metrics could be derived. Brain modularity, nodal strength and nodal diversity were determined for each participant and compared between all participants in the MRI cohorts.*

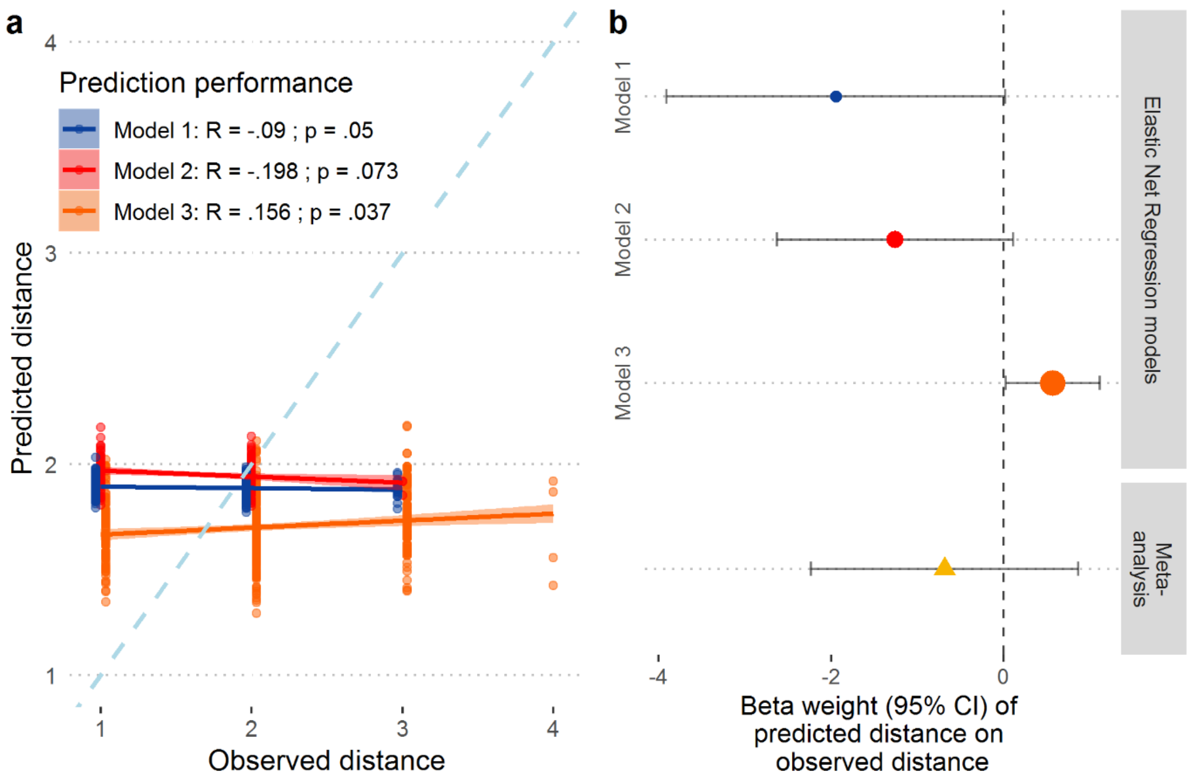


*Figure S1. Performance of regression models predicting social distance from similarity in whole-brain functional connectivity. a) Predicted vs. observed social distance in testing data sets for models 1, 2 and 3. Blue dotted line represents the line of perfect accuracy (predicted = observed). Pearson's correlations were negative for models 1 and 2, indicating that the models performed worse than chance. P values are from LME with crossed random effects, taking into account clustering of dyad members. b) Meta-analysis of LME models for regression model of predictive performance.*


1 Parkinson, C., Kleinbaum, A. M. & Wheatley, T. Similar neural responses predict friendship. *Nature Communications* **9**, 332, doi:10.1038/s41467-017-02722-7 (2018).

2 Stevens, S. S. Issues in psychophysical measurement. *Psychol. Rev.* **78**, 426 (1971).

3 Clauset, A., Newman, M. E. & Moore, C. Finding community structure in very large networks. *Physical review E* **70**, 066111 (2004).

4 Brant-Zawadzki, M., Gillan, G. D. & Nitz, W. R. MP RAGE: a three-dimensional, T1-weighted, gradient-echo sequence--initial experience in the brain. *Radiology* **182**, 769-775, doi:10.1148/radiology.182.3.1535892 (1992).

5 Woolrich, M. W., Ripley, B. D., Brady, M. & Smith, S. M. Temporal autocorrelation in univariate linear modeling of FMRI data. *Neuroimage* **14**, 1370-1386 (2001).

6 Jenkinson, M., Beckmann, C. F., Behrens, T. E., Woolrich, M. W. & Smith, S. M. FSL. *Neuroimage* **62**, 782-790 (2012).

7 Smith, S. M. *et al.* Advances in functional and structural MR image analysis and implementation as FSL. *Neuroimage* **23**, S208-S219 (2004).

8 Jenkinson, M. & Smith, S. A global optimisation method for robust affine registration of brain images. *Med. Image Anal.* **5**, 143-156 (2001).

9 Jenkinson, M., Bannister, P., Brady, M. & Smith, S. Improved optimization for the robust and accurate linear registration and motion correction of brain images. *Neuroimage* **17**, 825-841 (2002).

10 Andersson, J. L. R., Jenkinson, M. & Smith, S. TR07JA1 : Non-linear optimisation. (FMRIB Centre, Oxford, United Kingdom, 2007).
11 Andersson, J. L. R., Jenkinson, M. & Smith, S. TR07JA2 : Non-linear registration, aka Spatial normalisation. (FMRIB Centre, Oxford, United Kingdom, 2007).
12 Smith, S. M. Fast robust automated brain extraction. *Hum. Brain Mapp.* **17**, 143-155 (2002).
13 Beckmann, C. F. & Smith, S. M. Probabilistic independent component analysis for functional magnetic resonance imaging. *IEEE Trans. Med. Imaging* **23**, 137-152, doi:10.1109/tmi.2003.822821 (2004).
14 Griffanti, L. *et al.* ICA-based artefact removal and accelerated fMRI acquisition for improved resting state network imaging. *Neuroimage* **95**, 232-247, doi:http://dx.doi.org/10.1016/j.neuroimage.2014.03.034 (2014).
15 Salimi-Khorshidi, G. *et al.* Automatic denoising of functional MRI data: combining independent component analysis and hierarchical fusion of classifiers. *Neuroimage* **90**, 449-468, doi:10.1016/j.neuroimage.2013.11.046 (2014).
16 Smith, S. M. *et al.* Correspondence of the brain's functional architecture during activation and rest. *Proc. Natl. Acad. Sci. U. S. A.* **106**, 13040-13045 (2009).
17 Yarkoni, T., Poldrack, R. A., Nichols, T. E., Van Essen, D. C. & Wager, T. D. Large-scale automated synthesis of human functional neuroimaging data. *Nature methods* **8**, 665 (2011).
18 Menon, V. in *Brain mapping: An encyclopedic reference* (ed Arthur W Toga) (Academic Press, 2015).
19 Grayson, D. S. & Fair, D. A. Development of large-scale functional networks from birth to adulthood: A guide to the neuroimaging literature. *Neuroimage* **160**, 15-31, doi:10.1016/j.neuroimage.2017.01.079 (2017).
20 Barr, D. J., Levy, R., Scheepers, C. & Tily, H. J. Random effects structure for confirmatory hypothesis testing: Keep it maximal. *Journal of memory and language* **68**, 10.1016/j.jml.2012.1011.1001, doi:10.1016/j.jml.2012.11.001 (2013).
21 Murayama, K., Sakaki, M., Yan, V. X. & Smith, G. M. Type I error inflation in the traditional by-participant analysis to metamemory accuracy: a generalized mixed-effects model perspective. *J. Exp. Psychol. Learn. Mem. Cogn.* **40**, 1287-1306, doi:10.1037/a0036914 (2014).
22 Sirimongkolkasem, T. & Drikvandi, R. On Regularisation Methods for Analysis of High Dimensional Data. *Annals of Data Science*, 1-27 (2019).
23 James, G., Witten, D., Hastie, T. & Tibshirani, R. *An introduction to statistical learning*. Vol. 112 (Springer, 2013).
24 Friedman, J. H., Hastie, T. & Tibshirani, R. Regularization Paths for Generalized Linear Models via Coordinate Descent. *Journal of Statistical Software* **33**, 22, doi:10.18637/jss.v033.i01 (2010).
